# Discoidins are cytosolic lectins that recognize methylated L-rhamnose epitopes on mycobacterial glycolipids and glycopeptidolipids upon vacuole damage

**DOI:** 10.64898/2026.06.18.733164

**Authors:** Davide D’Amico, Gregory Goudge, Mélanie Foulon, Lourriel S. Macale, Larry S. Lico, Cheng-Kai Chen, Julien Prados, Ruixiang Blake Zheng, Aaron Franklin, Giselle Hevia Hernandez, Stephan von Gunten, Todd L. Lowary, Patrick Moynihan, Thierry Soldati

## Abstract

Mycobacteria are surrounded by a dynamic, glycolipid-rich envelope that governs virulence and immune recognition, yet how cytosolic lectins sense its organization during intracellular infection remains unclear. Here, we show that *Dictyostelium discoideum* discoidins act as cytosolic sensors of mycobacterial envelope exposure. During infection, *Mycobacterium marinum* damages the limiting vacuolar membrane, triggering recruitment of all canonical isoforms (DscA, DscC, DscD, and DscE) to distinct foci around bacilli. Discoidins are not recruited after sterile endolysosomal damage, indicating that, unlike galectins, they recognize non-self surface-associated bacterial glycans rather than exposed host glycans. DscA and DscE display distinct glycan specificities. Combining transposon mutagenesis, synthetic glycan arrays, biochemical fractionation, and envelope mutants, we identify a restricted set of lipid-associated determinants enriched in methylated L-rhamnose motifs within LOS-, PGL-, and GPL-related structures. Recognition requires the conserved β-galactoside-binding pocket of the H-type lectin domain, and isoform-specific synthetic disaccharides inhibit binding to intact bacteria. Analyses exclude AG–PG, AM/LAM, α-glucan, and TDM as major ligands and identify LOS and GPL-like glycolipids as dominant targets. PDIM/PGL or LOS disruption reduces discoidin binding to intact bacteria, although some ligands remain detectable in envelope extracts, underscoring the importance of accessibility. LOS deficiency also reduces DscA recruitment to intracellular bacilli. Therefore, binding of discoidins to glycan epitopes depends on ligand presence and accessibility, as well as on envelope perturbations and unmasking of specific glycolipids and glycopeptidolipids. These findings reveal pathogen envelope glycans as direct targets of cytosolic lectin surveillance and establish discoidins as probes of mycobacterial envelope remodeling during infection.

**Importance:** Many disease-causing bacteria are protected by an outer layer that promotes survival inside host cells. This layer is remodeled during infection, changing which molecules are exposed to the host, but how host cells detect this process remains unclear. Using the amoeba *Dictyostelium discoideum*, which shares defense mechanisms with human immune cells that engulf microbes, we show that carbohydrate-binding proteins called discoidins accumulate on intracellular *Mycobacterium marinum*, a close relative of *Mycobacterium tuberculosis*, which causes tuberculosis. Discoidins recognize specific sugar-containing molecules at the bacterial surface, but their recruitment depends not only on whether these targets are present but also on how they are displayed. Changes in envelope organization can therefore mask or expose discoidin targets. Our findings establish discoidins as sensors of bacterial surface remodeling and reveal an ancient surveillance mechanism through which host cells monitor changes in pathogen surface sugars.

## Introduction

Mycobacteria possess a specialized cell envelope that is structurally and biochemically distinct from those of classical Gram-positive and Gram-negative bacteria and is a major determinant of the pathogenicity of species such as *Mycobacterium tuberculosis* (Mtb) and other members of the Mtb complex (1). Broadly conserved across pathogenic mycobacteria, including non-tuberculous mycobacteria (NTM) such as *Mycobacterium marinum* (Mmar), this multilayered envelope comprises a plasma membrane; an arabinogalactan–peptidoglycan (AG–PG) scaffold covalently linked to long-chain mycolic acids; an unusual outer membrane known as the mycomembrane; and an extracellular capsule (2, 3). Beyond serving as a permeability barrier, the envelope is a dynamic and highly organized host– pathogen interface whose composition and biophysical properties influence intracellular adaptation, immune evasion, and virulence (1).

Although its core architecture is conserved, the outer envelope contains a diverse repertoire of extractable lipids and glycolipids that varies among species and growth states, including phenolic glycolipids (PGLs), phthiocerol dimycocerosates (PDIMs), sulfolipids, lipooligosaccharides (LOSs), and D-amino acid-containing glycopeptidolipids (GPLs). These molecules collectively contribute to environmental resistance and permeability control and shape the spatial organization and physicochemical properties of the bacterial surface (1, 4, 5). By controlling the exposure or masking of underlying determinants, they influence membrane organization, molecular accessibility, and host recognition (1). Many of these virulence-associated lipids are conserved between Mtb and Mmar, supporting Mmar as a powerful model for conserved mechanisms of mycobacterial pathogenesis.

Several envelope-associated lipids modulate host responses during infection (4, 5). PGL and sulfolipids can dampen macrophage inflammation by interfering with TLR-dependent signaling, including TLR2 antagonism by sulfoglycolipids and suppression of TLR4-TRIF signaling by PGL (6–8), whereas PDIM can spread into host membranes and alter their organization and biophysical properties (9, 10). Together with membrane-damaging secreted peptidic effectors, these lipids contribute to phagosomal disruption and cytosolic access during intracellular infection (11–13). Beyond these individual activities, the biological effects of mycobacterial glycolipids depend on their spatial distribution, membrane context, and surface accessibility, all of which can vary dynamically during bacterial growth and intracellular adaptation (1). Despite extensive characterization of individual lipids, considerably less is known about how envelope organization and remodeling regulate the surface exposure of glycolipid-associated ligands and, consequently, how these changes influence host recognition mechanisms during infection.

After phagocytosis, pathogenic mycobacteria establish a *Mycobacterium*-containing vacuole (MCV) whose maturation is actively manipulated to avoid lysosomal killing and digestion (14, 15). As infection progresses, effector molecules secreted through the ESX-1 type VII secretion system (T7SS), including the main membranolytic toxin EsxA, damage the MCV membrane and promote access to the host cytosol, a key step enabling further intracellular proliferation and dissemination (16–19).

Host recognition of extracellular mycobacteria relies heavily on glycan-binding receptors such as C-type lectin receptors (CLRs). The CLR DC-SIGN (dendritic cell-specific intercellular adhesion molecule 3-grabbing nonintegrin) recognizes lipoarabinomannan (LAM) and discriminates among mycobacterial species through selective recognition of the mannose caps of LAM (20–22). Nanoscale clustering of bacterial surface ligands also shapes pathogen–host interactions: ligand organization at the bacterial surface and clustering of host receptors both influence recognition (23). In the cytosol, soluble β-galactoside-binding galectins accumulate at damaged pathogen-containing vacuoles, where they sense host glycans aberrantly exposed to the cytosol by membrane damage (24). In addition to this intracellular role in membrane-damage surveillance, galectins can be secreted through unconventional pathways and act as soluble pattern recognition receptors (PRRs) for viruses, fungi, protists, helminths, and bacteria (25–29).

Several studies propose that galectins may also interact directly with mycobacterial surface glycoconjugates exposed after MCV damage. Although defined by their affinity for β-galactoside-containing glycans, *in vitro* studies suggest that their recognition repertoire may extend to structurally distinct mycobacterial glycoconjugates. Recombinant galectin-3 binds lipid species resembling phosphatidylinositol mannosides (PIMs) from *Mycobacterium bovis* BCG (30) and mycolic acids from Mtb (31). Galectin-9 also directly binds chemically synthesized Mtb arabinogalactan (AG), and this interaction aggravates infection outcomes in both Mtb-infected severe combined immunodeficient (SCID) mice and Mmar-infected zebrafish (32). However, these findings come mainly from extracellular bacteria or *in vitro* systems; direct interactions between galectins and intracellular mycobacteria after MCV damage have not been visualized during intracellular infection.

Discoidins are cytosolic lectins from *Dictyostelium discoideum* (*D. discoideum*) and were among the first lectins identified outside the plant kingdom (33). The *D. discoideum* genome encodes four classical isoforms that segregate into two subfamilies: Discoidin I comprises DscA, DscC, and DscD, whereas Discoidin II is represented by DscE. BLAST and domain architecture analyses also predict several discoidin-like proteins of unknown function. Among them, three proteins, Galactose-binding domain-containing protein (Gbdcp), DD7-1, and Discoidin-like protein retain the same bipartite organization found in canonical discoidins, consisting of an N-terminal discoidin domain and a C-terminal H-type lectin domain. Group I discoidins share approximately 97% sequence identity, whereas DscE has only ∼47% identity to group I members. Structural studies show that discoidins form trimers whose β-galactoside-binding site lies in a shallow groove at the interface between adjacent H-type lectin domains (34). Despite their distinct evolutionary origins and the lack of sequence or structural homology, discoidins and galectins show functional convergence in cytosolic localization, β-galactoside binding, and unconventional secretion. Early studies showed that discoidins bind bacterial lysates and fixed *Klebsiella pneumoniae* and *Escherichia coli* (35, 36). Proteomic analysis further revealed selective enrichment of DscA and DscE in compartments containing virulent *Legionella pneumophila* but not avirulent species *L. hackeliae* (37), suggesting context-specific engagement in pathogen recognition. We recently identified discoidins as cytosolic lectins that accumulate at damaged MCVs in a T7SS-dependent manner and selectively recognize exposed bacterial glycans. Recruitment required ligand accessibility during vacuole rupture, and loss of discoidins altered bacterial access to the cytosol and intracellular fate, supporting a role for these lectins in cell-autonomous immunity (38).

Here, we combine live-cell imaging, genome-wide transposon library screening, synthetic glycan arrays, and biochemical fractionation to identify Mmar surface ligands required for discoidin binding. We show that discoidins selectively recognize a restricted set of lipid-linked glycoconjugates enriched in methylated L-rhamnose residues, including LOS-, GPL-, and PGL-associated structures. Binding depends on PDIM- and LOS-mediated surface organization: removing these lipids markedly reduces lectin binding to live bacteria even when candidate ligands remain biochemically detectable by dot blot, showing that glycan presentation in the outermost mycobacterial layer enables discoidin recognition. These findings establish discoidins as intracellular probes of mycobacterial surface organization and dynamics, providing a route toward direct visualization and turnover during infection.

## Results

### Discoidins are recruited to intracellular mycobacteria and selectively recognize non-self glycans

To investigate the cellular localization of the four classical discoidin isoforms during mycobacterial infection, we examined live *D. discoideum* cells expressing GFP-DscA, GFP-DscC, GFP-DscD or GFP-DscE and infected with Mmar expressing cytoplasmic mCherry. Fluorescence microscopy revealed that all four isoforms formed discrete foci associated with intracellular bacilli (Figure 1A).

**Figure 1.**
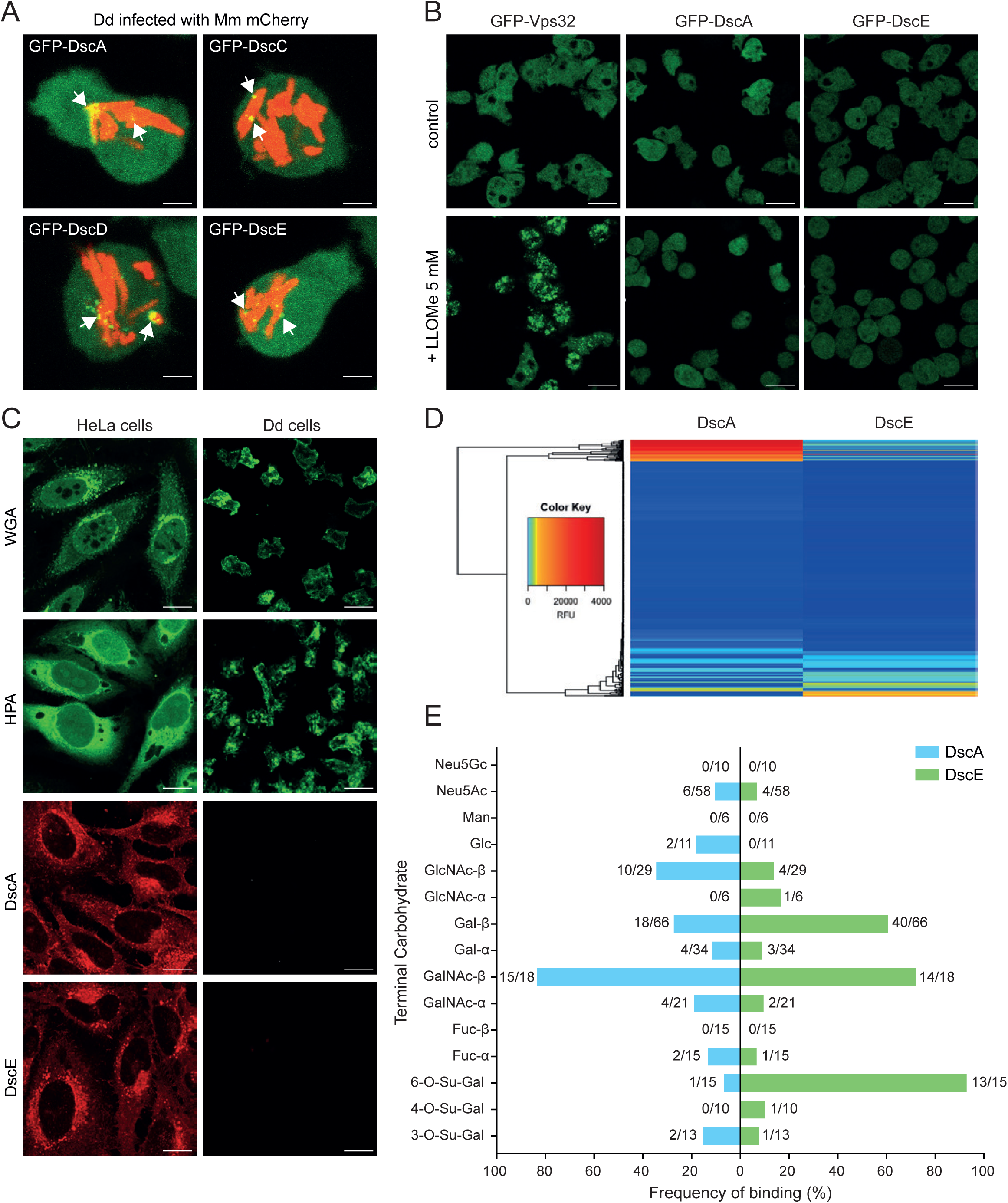
Discoidins recognize non-self glycans and display distinct glycan-binding profiles. **(A)** Representative live confocal images of *D. discoideum* cells expressing GFP-DscA, GFP-DscC, GFP-DscD or GFP-DscE (green) infected with mCherry-expressing Mmar (red). All four canonical discoidin isoforms form discrete foci associated with intracellular bacilli (arrows). Scale bars, 2 µm. **(B)** Images from live-cell confocal time-lapse microscopy of *D. discoideum* cells expressing GFP-Vps32, GFP-DscA or GFP-DscE, shown at 0 and 10 min after addition of 5 mM LLOMe. GFP-Vps32, an ESCRT-III component, was used as a positive control for the membrane repair response. Scale bars, 10 µm. **(C)** Fluorescence microscopy of fixed and permeabilized HeLa and *D. discoideum* cells incubated with AF488-labeled wheat germ agglutinin (WGA) or *Helix pomatia* agglutinin (HPA) (green), or with CF640R-labeled recombinant DscA or DscE (red). Scale bars, 10 µm. **(D)** Heat map of DscA and DscE binding to an expanded mammalian and bacterial glycan array. Rows represent individual glycans hierarchically clustered according to their binding profiles; colors indicate background-subtracted binding intensity expressed as relative fluorescence units (RFU). **(E)** Frequency of DscA (blue) and DscE (green) binding to glycans grouped according to their terminal carbohydrate. Numbers indicate the number of bound glycans relative to the total number of glycans within each category.

Structural comparison of the trimeric structures of DscA and DscE obtained after X-ray crystallography (34) with the AlphaFold-predicted models of DscC and DscD revealed only minor differences among DscA, DscC and DscD, whereas DscE differed more extensively, including within the β-galactoside binding pocket (Suppl. Figure S1A). An amino acid sequence identity matrix further showed 95.3–97.6% pairwise identity among DscA, DscC and DscD, while DscE shared only 47.0– 47.4% identity with the three Discoidin I isoforms (Suppl. Figure S1B). Accordingly, subsequent experiments focused on DscA as a representative of the highly conserved discoidin I family and DscE as the sole discoidin II isoform.

In animal cells, cytosolic galectins are known to bind self-glycans exposed upon sterile endolysosomal membrane damage (39). To determine whether this also occurs with discoidins, *D. discoideum* cells expressing GFP-DscA or GFP-DscE were treated with 5 mM L-leucyl-L-leucine methyl ester (LLOMe), a lysosomotropic compound that induces endolysosomal membrane permeabilization (40–43). GFP-Vps32, an ESCRT-III component was used as a positive control for the membrane repair response. LLOMe treatment induced a pronounced redistribution of GFP-Vps32 into punctate structures, whereas GFP-DscA and GFP-DscE remained diffusely cytosolic (Figure 1B). Thus, LLOMe-induced endolysosomal damage did not reproduce the discoidin localization observed during Mmar infection.

To confirm if discoidins bind glycans present in growing, non-differentiating, *D. discoideum* cells, purified recombinant and fluorescently labeled CF640R-DscA and CF640R-DscE were incubated with fixed and permeabilized cells and HeLa cells were used as a heterologous reference. AlexaFluor488 (AF488)-labeled wheat germ agglutinin (WGA) and *Helix pomatia* agglutinin (HPA) were also used as control lectins. Both WGA and HPA bound HeLa and *D. discoideum* cells, confirming the presence of accessible and conserved glycans (Figure 1C). CF640R-DscA and CF640R-DscE also labeled HeLa cells but showed no appreciable binding to *D. discoideum* cells (Figure 1C).

We next profiled the glycan-binding specificities of DscA and DscE using an expanded mammalian and bacterial glycan array. Analysis of the array data identified clusters of glycans both common and specific to DscA and DscE (Figure 1D). Consistent with previous studies (34), both isoforms recognized most β-GalNAc-terminated glycans represented on the array (15/18 for DscA and 14/18 for DscE). DscE also frequently recognized glycans terminating in β-Gal (40/66) or 6-*O*-sulfated Gal (13/15), whereas these structures were less frequently bound by DscA (18/66 and 1/15, respectively) (Figure 1E). Together, these results show that DscA and DscE do not detectably bind growing *D. discoideum* cells, which is consistent with recent reports showing that they do not synthesize glycans with β-galactoside epitopes. Moreover, these data indicate that, in addition to β-galactosides, DscA and DscE recognize a broader repertoire of glycan epitopes.

## Cytosolic discoidins directly bind the mycobacterial surface

To examine in greater detail how discoidins associate with intracellular mycobacteria, we performed three-dimensional high-resolution reconstructions of Mmar-infected *D. discoideum* cells expressing GFP-DscA or GFP-DscE. Both isoforms formed discrete foci and patches decorating the surface of bacilli following MCV damage and bacteria escape to the cytosol, while additional discoidin puncta were frequently observed at a distance from intracellular bacteria (Figure 2A).

**Figure 2.**
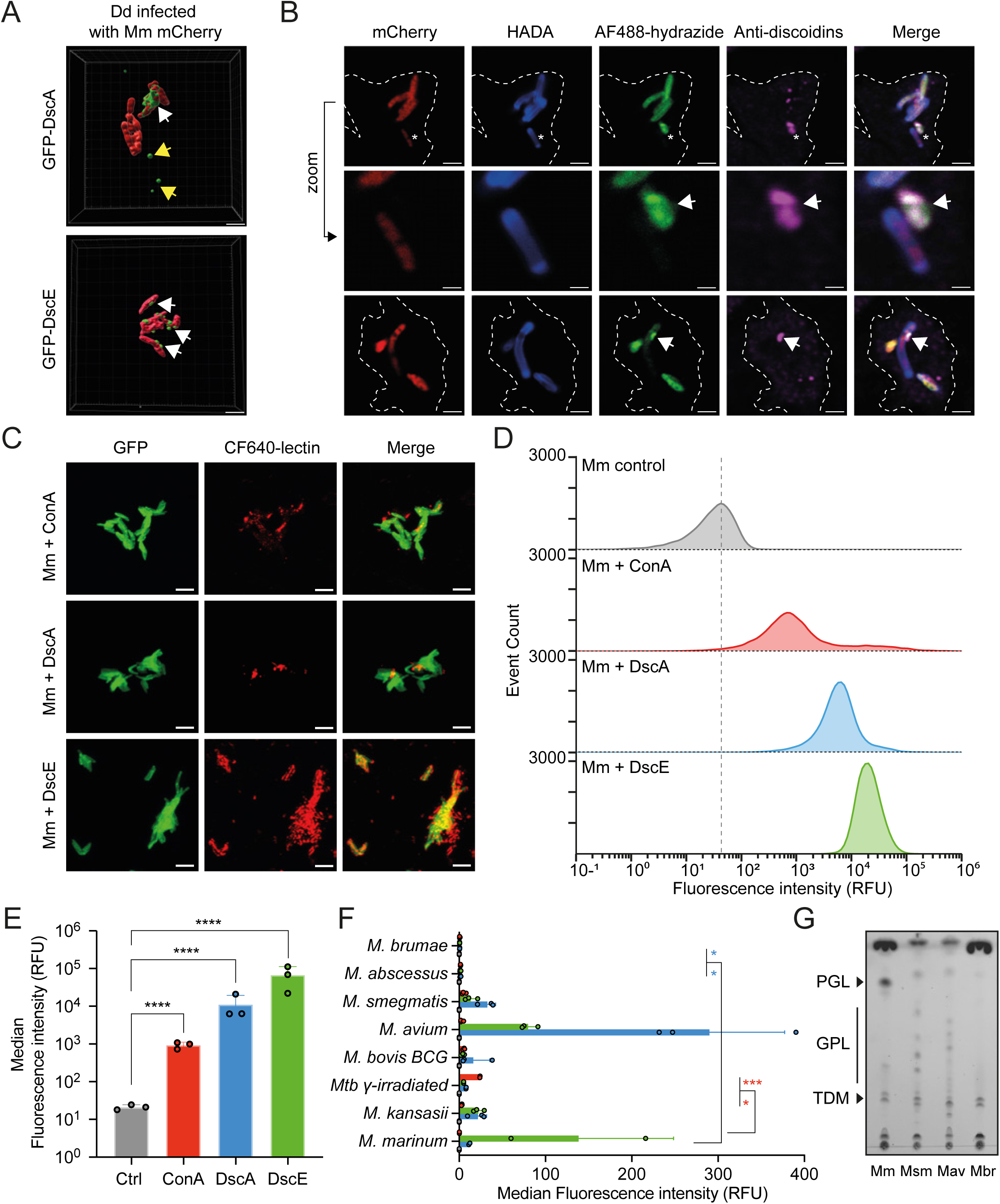
Discoidins bind mycobacterial surfaces and reveal heterogeneous glycan exposure. **(A)** Representative 3D reconstructions of Mmar (red) in *D. discoideum* cells expressing GFP-DscA (top) or GFP-DscE (bottom). White arrows indicate discrete discoidin foci associated with bacterial surfaces, whereas yellow arrows indicate additional cytosolic puncta. **(B)** Labeling of intracellular Mmar expressing cytoplasmic mCherry (red), stained with HADA (blue) to label sites of active peptidoglycan synthesis and with AF488-hydrazide (green) to label oxidized carbohydrates. Immunostaining with an antibody recognizing Discoidin I and II (magenta) reveals localized patches of discoidins at the bacterial surface (arrowheads), partially overlapping with hydrazide-positive regions. Dashed lines indicate host cell boundaries. Upper and lower panels, scale bars, 5 µm; zoomed views, scale bars, 1 µm. **(C)** Representative fluorescence images of Mmar expressing cytoplasmic GFP (green) incubated with CF640R-labeled lectins (red): ConA, DscA or DscE. Scale bars, 5 µm. **(D)** Flow cytometry analysis of lectin binding to Mmar. Histograms show fluorescence intensity distributions for control bacteria (gray), ConA (red), DscA (blue) and DscE (green). **(E)** Quantification of lectin binding shown as median fluorescence intensity. Individual points represent three independent experiments. Statistical significance was assessed by one-way ANOVA followed by Dunnett’s multiple-comparisons test, comparing each lectin with the control. **(F)** Flow cytometry analysis of discoidin binding across mycobacterial species. Median fluorescence intensity is shown for control (gray), DscA (blue), DscE (green) and ConA (red). Individual points represent three independent experiments. Statistical significance was assessed by two-way ANOVA followed by uncorrected Fisher’s LSD test. Pairwise comparisons are indicated by brackets. **(G)** Thin-layer chromatography analysis of lipid extracts from selected mycobacterial species. Major glycolipid classes (PGL, GPL and TDM) are indicated. Mm, *M. marinum*; Msm, *M. smegmatis*; Mav, *M. avium*; Mbr, *M. brumae*. *P < 0.05, ***P < 0.001 and ****P < 0.0001. Dd, *D. discoideum*; Mm, *M. marinum*.

To better decipher what is recognized on the bacterial surface, various envelope components were fluorescently labeled. Mmar surface-exposed glycoconjugates were selectively labeled by mild periodate treatment followed by AF488-hydrazide coupling. Mmar were visualized by constitutive cytoplasmic mCherry expression together with metabolic incorporation of HADA (3-[[(7-Hydroxy-2-oxo-2*H*-1-benzopyran-3-yl)carbonyl]amino]-D-alanine hydrochloride) into the nascent peptidoglycan (Figure 2B). Immunostaining for endogenous Discoidin I and II revealed co-localization of discoidin foci with the bacterial surface, some co-localizing with patches of AF488-hydrazide labeled components at the surface and at some distance from the intracellular bacilli (Figure 2B).

To validate that discoidin recruitment reflects direct binding to bacterial surface components, we next performed *in vitro* labeling assays using recombinant fluorescently labeled CF640R-DscA and CF640R-DscE. Incubation with broth-grown Mmar resulted in surface association visualized by fluorescence microscopy (Figure 2C). The binding pattern was comparable to that observed with Concanavalin A (ConA), a lectin known to recognize mannose-containing glycoconjugates on mycobacteria and especially man-LAM (44). Flow cytometric analysis confirmed direct binding, with both discoidins inducing a pronounced shift in fluorescence intensity relative to the negative control, and to ConA (Figure 2D,E). CF640R-DscE produced a stronger shift than CF640R-DscA, indicating isoform-specific differences in binding efficiency, likely reflecting the combined effects of ligand accessibility and availability, as well as differences in effective affinity and avidity within the native surface context.

We next evaluated discoidin binding across multiple mycobacteria species (Figure 2F). The relative intensity of DscA and DscE binding varied across species. In several cases DscA produced higher fluorescence signals, whereas DscE displayed the highest signal overall for Mmar. ConA also exhibited a distinct binding distribution across species. Overall, strong binding of both DscA and DscE was observed for Mmar and *Mycobacterium avium*. Intermediate binding profiles were detected for *Mycobacterium kansasii*, *M. bovis* BCG and *Mycobacterium smegmatis*, whereas *Mycobacterium abscessus* and *Mycobacterium brumae* showed minimal or near-background signal. γ-irradiated Mtb exhibited detectable but comparatively lower binding than Mmar and *M. avium*. To explore whether differential binding correlated with envelope composition, we compared the glycolipid profiles of representative high- and low-binding species by thin-layer chromatography (TLC) (Figure 2G). High-binding species exhibited a broader repertoire of surface-associated glycolipids, including phenolic glycolipids (PGLs) in Mmar and potential glycopeptidolipids (GPLs) in *M. avium* and *M. smegmatis*, whereas a low-binding species such as *M. brumae* showed no detectable glycolipid species. These findings suggest that discoidin recognition correlates with the presence of abundant glycan-bearing outer envelope glycolipids, rather than with a single conserved lipid class.

### Screening of an Mmar library of transposon mutants identifies key cell envelope-associated pathways required for discoidin binding

To define the bacterial determinants required for discoidin recognition, we performed a genome-wide screen using an Mmar transposon insertion library and FACS after incubation with CF640R-DscE, the isoform with the strongest binding signal (Figures 2D,E and 3A). Bacilli in the lowest 5% of the fluorescence distribution were isolated, enriching clones with markedly reduced binding. Because the initial distribution spanned almost two orders of magnitude, the sorted population was expanded and subjected to two further rounds of labeling and enrichment to distinguish stable mutants from transient phenotypic variants. The low-fluorescence fraction progressively expanded across rounds (Figure 3B). Deep sequencing mapped transposon insertions in the input and enriched populations, and TA-site abundance was compared between rounds (Figure 3C). Volcano plots identified 43, 104 and 323 significantly overrepresented TA sites after rounds 1, 2 and 3, respectively, alongside a broader set of depleted insertions. Sites enriched in all three rounds were considered high-confidence candidates whose disruption reduces surface binding. In total, 42 genes contained significant TA insertions. Cell-envelope functions were strongly overrepresented (Figure 3D; Table S1): 28.5% of hits were linked to the cell wall or its biosynthesis and 14.3% to T7SS, a secretion system related to the cell envelope. The latter group included five likely secreted proteins: a putative toxin (MMAR_0480) and adjacent TAP/Esx partners (MMAR_0481 and MMAR_0482), a PPE protein (MMAR_1129), and a PE-PGRS protein (MMAR_2102). T7SS-secreted proteins can shape cell-surface characteristics, including capsular integrity (45). Other hits involved central metabolism, regulatory systems and DNA/RNA pathways. Several enriched loci, including *fadD2*, *fadD28* and *mas*, participate in PDIM/PGL biosynthesis. These findings link DscE binding to genetically encoded pathways controlling outer-envelope lipid biosynthesis and organization. The enrichment of PDIM/PGL biosynthetic genes suggests that this pathway contributes to discoidin binding, either directly or by modulating ligand accessibility at the bacterial surface.

**Figure 3.**
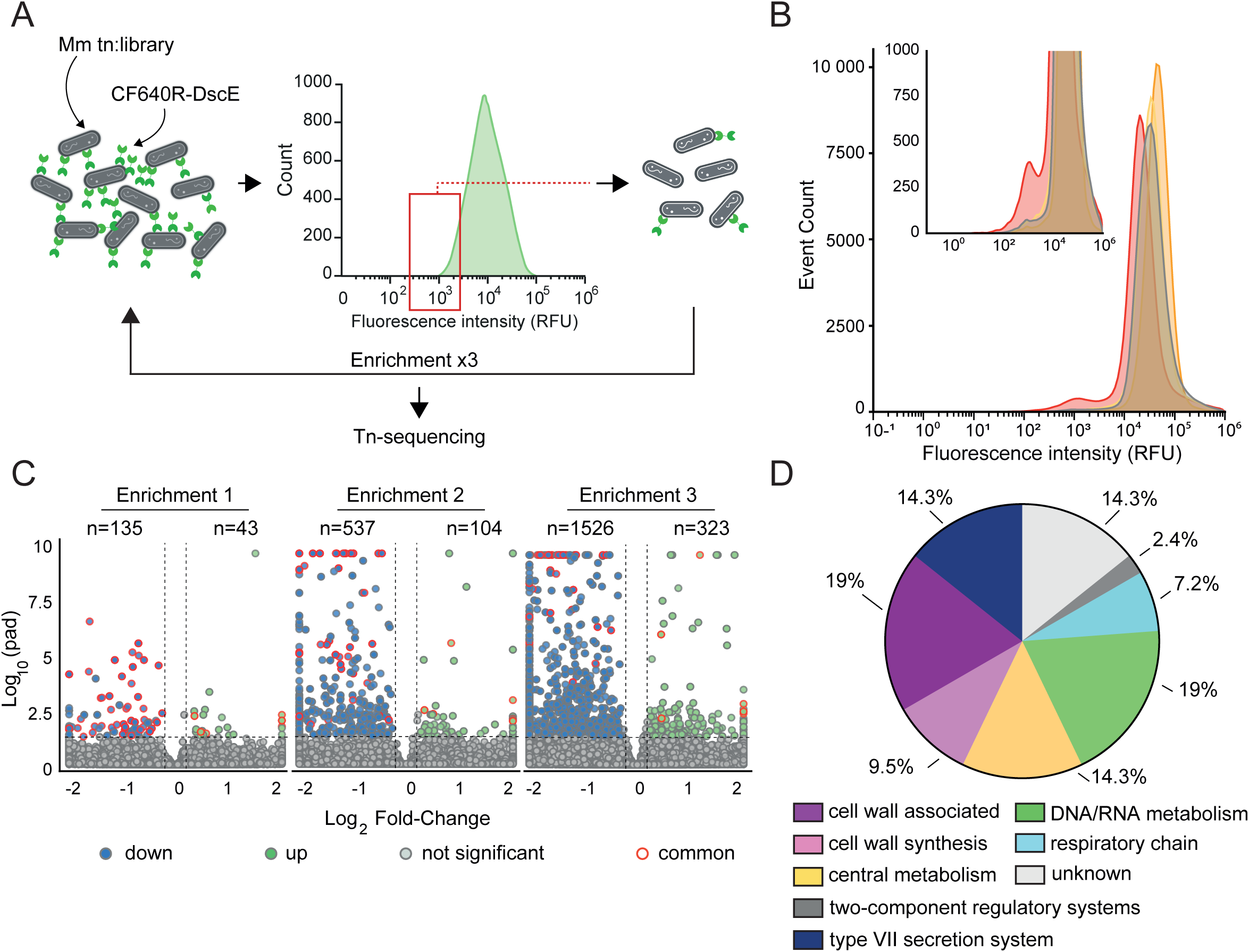
Transposon-based screen identifies determinants of discoidin binding in Mmar. **(A)** Schematic overview of the screening strategy. An Mmar transposon mutant library was incubated with CF640R-labeled DscE and subjected to FACS sorting to isolate the lowest 5% fluorescence population. Sorted bacteria were expanded and re-sorted for three consecutive rounds prior to transposon sequencing. **(B)** Flow cytometry analysis of DscE binding during enrichment. Histograms show fluorescence intensity distributions for the initial library and after each round of enrichment (gray indicates the unsorted input population, yellow the first low-binding enrichment round, orange the second enrichment round, and red the third sorting round). The inset shows a magnified view of the low-fluorescence region used for sorting. **(C)** Volcano plots showing differential representation of transposon insertions after each enrichment round. Log₂ fold-change is plotted against –log₁₀ (adjusted p-value). Enriched insertions are shown in green, depleted insertions in blue, non-significant in gray, and insertions shared across enrichment rounds in red. **(D**) Functional classification of genes enriched in the low-binding population after the third round of selection. Categories are shown based on annotated pathways.

### Discoidin binding is detected in multiple mycobacterial envelope fractions

To dissect the envelope components contributing to binding, the Mmar envelope was fractionated into capsular, polar-lipid and apolar-lipid pools (Figure 4A,B). Both DscA and DscE bound all three fractions by dot blot, with stronger signals in the capsular and polar-lipid pools (Figure 4B). This pattern suggested recognition of multiple envelope species carrying shared or similar terminal sugar motifs. Because these fractions may contain both loosely associated surface material and components from deeper envelope layers, we next tested defined structural elements. The carbohydrate-rich arabinogalactan– peptidoglycan (AG–PG) matrix was a plausible candidate: synthetic mycobacterial arabinogalactan interacts with galectin-9 *in vitro*, and galectin-9 binds intact Mtb H37Rv, *M. bovis* BCG and *M. smegmatis* (32). Quantitative pull-down assays used purified insoluble Mmar AG–PG and rhodamine-WGA as a positive control for accessible N-acetylglucosamine-containing structures. WGA was retained by both 150 and 750 µg AG–PG, with an approximately two-fold stronger signal at 750 µg, confirming ligand availability in the preparation (Figure 4C). Under the same conditions, CF640R-DscA and CF640R-DscE showed no significant association with 150 µg AG–PG (Figure 4D,E, left panels; Suppl. Figure S2). Repeating the assay with 750 µg did not change this result: discoidin signals remained almost entirely in the unbound and wash fractions (Figure 4D,E, right panels), indicating negligible interaction with the AG–PG. These results exclude the AG–PG as a primary discoidin ligand.

**Figure 4.**
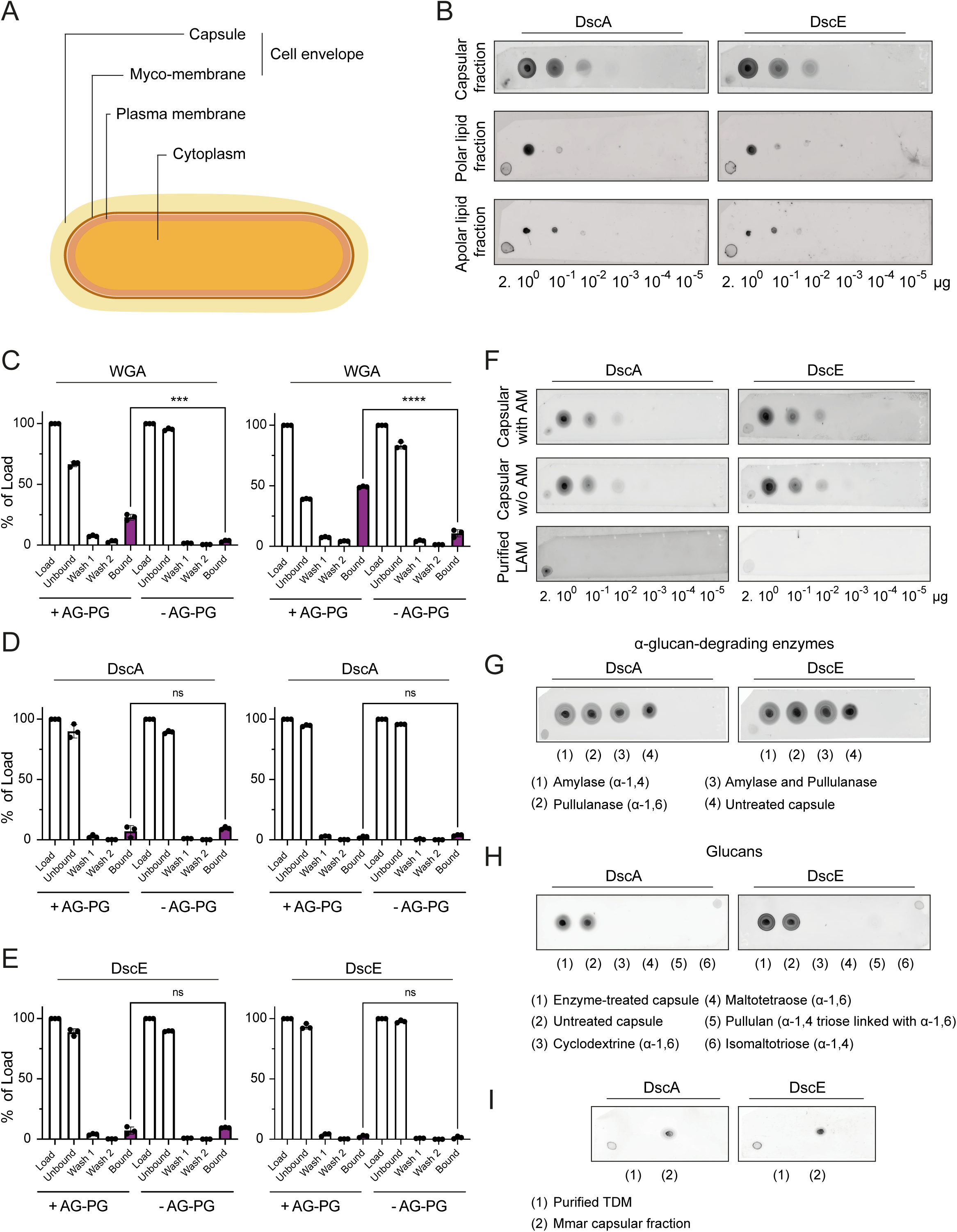
Discoidin binding is associated with capsular and polar lipid fractions and excludes major envelope glycans. **(A**) Schematic representation of the Mmar cell envelope, indicating the capsule, mycomembrane, plasma membrane and cytoplasm. **(B)** Dot blot analysis of Mmar envelope fractions probed with CF640R-labeled DscA or DscE. Capsular, polar lipid and apolar lipid fractions were spotted at decreasing concentrations. **(C)** Binding of rhodamine-labeled WGA to purified 150 µg (left) and 750 µg AG–PG (right) (+AG–PG) or AG-depleted samples (-AG–PG). Data are expressed as percentage of input. **(D-E)** Binding of CF640R-labeled DscA (D) and DscE (E) to purified 150 µg (left) and 750 µg AG– PG (right) (+AG–PG) or AG-depleted samples (−AG–PG). Data are expressed as percentage of input. For panels C–E, statistical significance was assessed using an unpaired two-tailed Student’s *t*-test comparing the bound fractions of +AG–PG and −AG–PG samples at each amount. ns, not significant; \*\*\**P* < 0.001; \*\*\*\**P* < 0.0001. **(F)** Dot blot analysis of capsular extracts with or without AM, and purified LAM, probed with CF640R-labeled DscA or DscE. **(G)** Dot blot analysis of capsular extracts following treatment with amylase (1), pullulanase (2), both enzymes (3), or no enzymatic treatment (4), probed with CF640R-labeled DscA or DscE. **(H)** Dot blot analysis of enzyme-treated capsule (1), untreated capsule (2), cyclodextrin (3), maltotetraose (4), pullulan (5) and isomaltotriose (6), probed with CF640R-labeled DscA or DscE. **(I)** Dot blot analysis of purified TDM (1) and the Mmar capsular fraction (2), probed with CF640R-labeled DscA or DscE. AG–PG, arabinogalactan–peptidoglycan; AM, arabinomannan; LAM, lipoarabinomannan; TDM, trehalose dimycolate; Mmar, *M. marinum*.

We next evaluated candidate capsular and apolar ligands. Discoidin binding was comparable between capsule extracts containing or depleted of arabinomannan (AM), and neither CF640R-DscA nor CF640R-DscE showed detectable binding to purified LAM spotted at 2 μg and in subsequent 10-fold serial dilutions (Figure 4F). These results exclude both AM and its lipid-anchored precursor LAM. Because α-glucan is a documented component of the Mmar capsule and a major mycobacterial capsular polysaccharide (46), extracts were treated with α-amylase and pullulanase, alone or together. Enzyme-treated samples retained signals indistinguishable from untreated controls (Figure 4G), excluding α-glucan as the primary ligand. Consistently, purified maltotetraose, isomaltotriose, pullulan and cyclodextrin failed to support detectable DscA or DscE binding (Figure 4H), showing that α-linked glucose polymers are not direct ligands. Finally, purified trehalose dimycolate, a major apolar virulence lipid, showed no signal relative to the Mmar apolar-lipid positive control (Figure 4I). Together, these experiments show that discoidins bind components present in several envelope fractions while excluding the AG–PG scaffold, AM, LAM, α-glucans and TDM as primary ligands.

### Discoidins selectively recognize lipid-associated mycobacterial glycans enriched in methylated L-rhamnose motifs

Commercial mammalian glycan arrays showed preferential discoidin binding to β-GalNAc-, glucose- and β-galactose-containing structures (34), but they did not contain mycobacterial glycans. We therefore used a previously developed synthetic array (47) comprising defined motifs from the major classes of mycobacterial surface carbohydrates (Suppl. Figures S3 and S4): LAM-related glycans (n = 29), PGL-related glycans (n = 18), LOS fragments (n = 4), GPLs (n = 3), a phosphatidyl-myo-inositol mannoside (n = 1), and capsular α-glucans (n = 6). The LOS motifs were derived from *M. kansasii*, whereas the PGL panel represented a broader range of mycobacterial glycolipids. Unlike arrays dominated by pyranose-containing glycans, this platform also included arabinofuranose-containing motifs. All glycans were chemically synthesized with defined linkages and stereochemistry and immobilized through BSA conjugation, allowing direct comparison of related structures. DscA and DscE selectively recognized a restricted subset of lipid-linked glycoconjugates (Figure 5A,B). Their profiles largely overlapped, although DscA showed slightly broader recognition and the isoforms differed modestly in signal intensity and glycan ranking (Suppl. Figure S3). Glycans reproducibly recognized by both isoforms clustered in specific lipid-associated classes (Figure 5C): PGL-related structures (IDs 30, 31, 36, 43 and 53), LOS fragments (IDs 55 and 62), and GPL-associated glycans (IDs 47 and 61). Despite their different architectures, most bound structures contained methylated L-rhamnose, often in terminal or subterminal positions. LAM-derived glycans and capsular α-glucans were not significantly recognized, consistent with the biochemical exclusions presented in Figure 4F-H.

**Figure 5.**
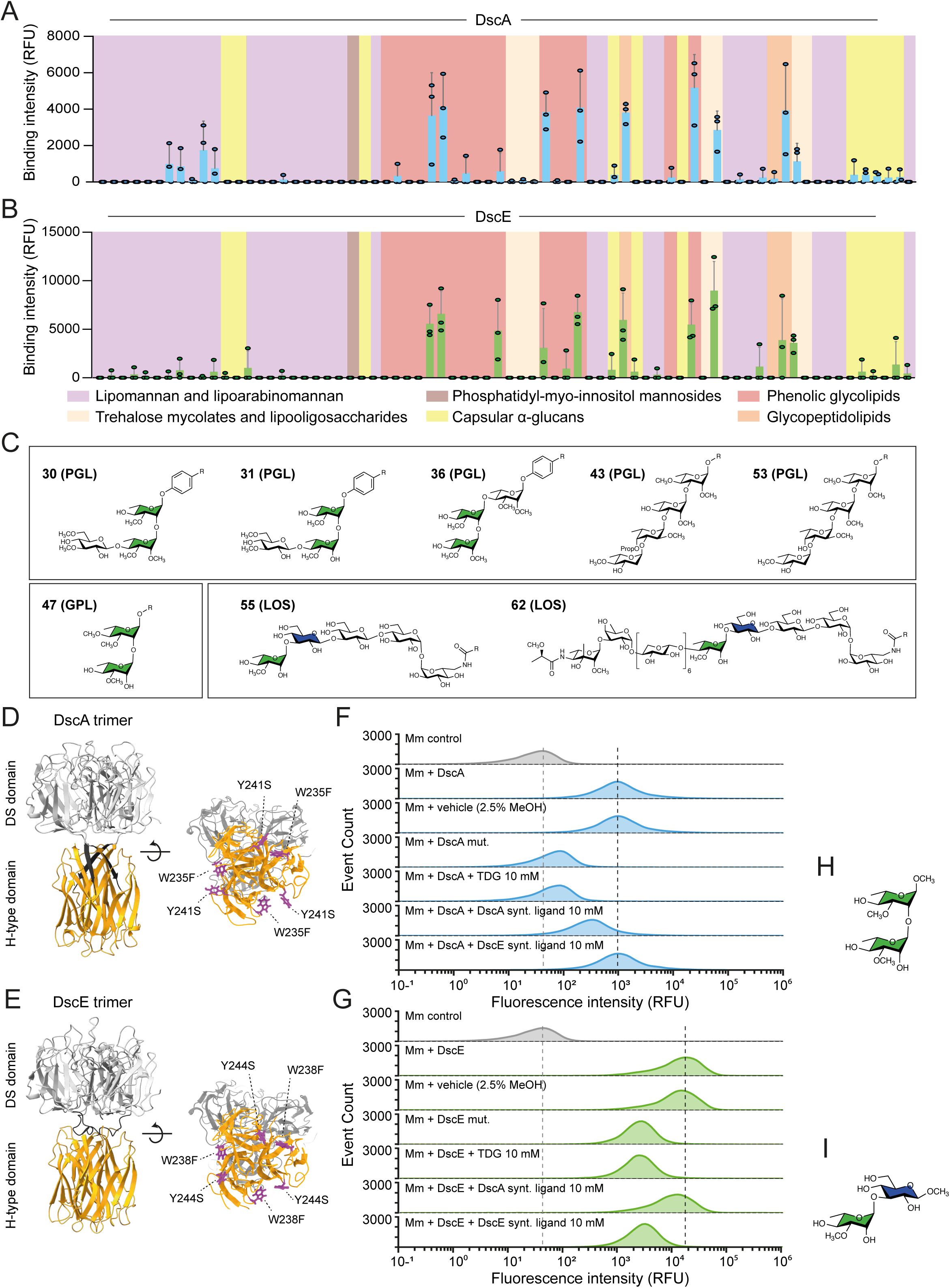
Discoidins selectively recognize lipid-associated mycobacterial glycans through the H-type lectin binding pocket and are differentially inhibited by synthetic ligands. **(A, B)** Glycan array analysis of CF640R-labeled DscA **(A)** and DscE **(B)** binding to synthetic mycobacterial glycans. Binding intensity (RFU) is shown for individual glycans grouped by class, including lipomannan and lipoarabinomannan, trehalose mycolates and lipooligosaccharides, phosphatidyl-*myo*-inositol mannosides, capsular α-glucans, phenolic glycolipids and glycopeptidolipids. **(C)** Representative structures of glycans showing reproducible discoidin binding, including PGL-related glycans (IDs 30, 31, 36, 43 and 53), a GPL-related glycan (ID 47) and LOS-related glycans (IDs 55 and 62). Highlighted residues indicate shared structural features among recognized glycans: methylated L-rhamnose (green) and glucose (blue). **(D, E)** Structural representations of the DscA **(D)** and DscE **(E)** trimers, showing the N-terminal discoidin (DS) domain (gray) and the C-terminal H-type lectin domain (orange). Mutations introduced into the lectin-binding pocket are indicated: W235F and Y241S in DscA, and W238F and Y244S in DscE. **(F, G)** Representative flow cytometry histograms of Mm incubated with CF640R-labeled DscA **(F)** or DscE **(G)**. Conditions include the wild-type lectin alone or in the presence of vehicle (2.5% MeOH), TDG, Ligand A or Ligand B (10 mM each), as well as the corresponding lectin-pocket mutant. Unlabeled Mm is shown as the control. Dashed lines indicate the fluorescence peaks of unlabeled bacteria and bacteria labeled with the corresponding wild-type lectin. **(H, I)** Chemical structures of Ligand A, methyl 2-*O*-(3-*O*-methyl-α-L-rhamnopyranosyl)-3-*O*-methyl-α-L-rhamnopyranoside **(H)**, and Ligand B, methyl 3-*O*-methyl-α-L-rhamnopyranosyl-(1→3)-β-D-glucopyranoside **(I)**. LM, lipomannan; LAM, lipoarabinomannan; LOS, lipooligosaccharide; PIM, phosphatidyl-*myo*-inositol mannoside; PGL, phenolic glycolipid; GPL, glycopeptidolipid; TDG, thiodigalactoside; Mm, *M. marinum*.

To determine whether recognition used the β-galactoside-binding pocket defined by the crystal structures (34), conserved aromatic residues in the H-type lectin domains were mutated in DscA and DscE (DscA W235F/Y241S; DscE W238F/Y244S) (Figure 5D,E). The DscA mutant showed reduced binding to several PGL- and LOS-related glycans but retained recognition of a subset of ligands (Suppl. Figure S3C). The DscE mutant lost signal from nearly all glycans recognized by the wild-type protein, particularly PGL- and GPL-associated structures (Suppl. Figure S3D). Binding to mycobacterial glycolipid glycans therefore largely requires an intact pocket, with subtle isoform-specific differences in ligand engagement. Flow cytometry after discoidin binding to intact bacteria confirmed this requirement: pocket mutations abolished DscA binding and reduced DscE binding ten-fold (Figure 5F,G). Thiodigalactoside (TDG), a soluble competitive inhibitor of cytosolic galectin β-galactoside binding (48), likewise fully suppressed DscA binding and reduced DscE binding ten-fold. Guided by the array profiles, two glycan epitopes were then synthesized as soluble competitors. Methyl 3-O-methyl-α-L-rhamnopyranosyl-(1→2)-3-O-methyl-α-L-rhamnopyranoside (DscA ligand) is an α-(1→2)-linked dirhamnoside carrying a 3-O-methyl group on both ʟ-rhamnose residues, whereas methyl 3-O-methyl-α-L-rhamnopyranosyl-(1→3)-β-D-glucopyranoside (DscE ligand) carries a single 3-O-methylated-L-rhamnose α-(1→3)-linked to a methyl β-D-glucopyranoside (Figure 5H, I). At 10 mM, the DscA ligand markedly reduced DscA binding while having little effect on DscE; conversely, the DscE ligand strongly reduced DscE binding without appreciably affecting DscA (Figure 5F,G). These reciprocal effects confirm that the distinct array-defined ligand preferences are reflected in discoidin binding to intact Mmar.

Together, DscA and DscE recognize a restricted set of PGL-, GPL- and LOS-associated glycans enriched in methylated L-rhamnose. Recognition requires an intact H-type lectin pocket, and isoform-selective methylated L-rhamnose disaccharides inhibit binding to intact Mmar.

### Discoidins recognize non-proteinaceous glycoconjugates enriched in the polar lipid fraction of *M. marinum*

To further characterize the biochemical nature of the discoidin-binding material present in capsular extracts, we assessed the sensitivity to proteolytic treatment. Capsular extracts were therefore treated with Proteinase K prior to dot blot analysis. In *M. bovis* BCG, protease treatment completely abolished discoidin binding (Suppl. Figure S5A). In contrast, treatment of Mmar capsular extracts reduced discoidin binding by approximately one order of magnitude but did not eliminate the signal (Figure 6A). These results indicate that the major discoidin ligand(s) of BCG is sensitive to Proteinase K, whereas a substantial fraction of the Mmar ligand(s) is protease resistant, consistent with glycolipid component such as GPLs, PGLs, LOSs or other related envelope-associated glycoconjugates.

**Figure 6.**
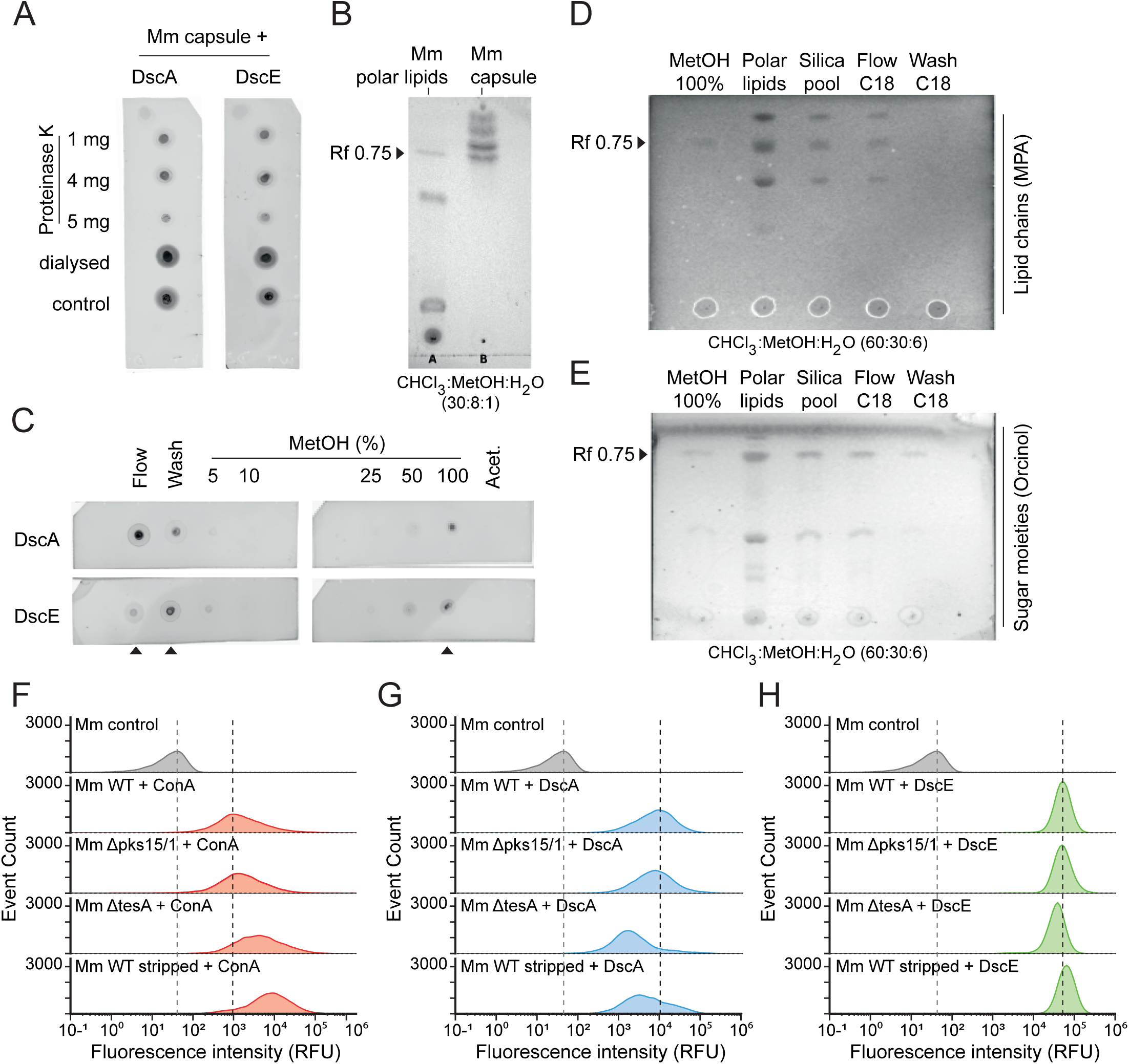
Discoidin binding is associated with a polar glycolipid and is modulated by mycobacterial envelope organization. **(A)** Dot blot analysis of 2-µL aliquots of Mm capsular extracts previously treated with proteinase K at concentrations of 1, 4 or 5 mg/mL, alongside dialyzed and untreated controls, and probed with CF640R-labeled DscA or DscE. **(B)** Thin-layer chromatography (TLC) of Mm polar lipid and capsular extracts using CHCl₃–MeOH–H₂O (30:8:1), revealing a shared band at *Rf* ≈ 0.75. **(C)** Dot blot analysis of polar lipid fractions obtained by C18 reverse-phase chromatography and probed with CF640R-labeled DscA or DscE. Fractions include the flow-through, wash, increasing MeOH concentrations (5–100%) and an 100% acetonitrile eluate. Binding activity was strongest in the flow-through and wash fractions and weaker in the 100% MeOH eluate; the acetonitrile eluate was negative. **(D)** Analytical TLC of the 100% MeOH eluate, unfractionated polar lipids, silica pool, C18 flow-through and C18 wash fractions, resolved using CHCl₃–MeOH–H₂O (60:30:6) and stained with molybdophosphoric acid (MPA) to detect lipids. **(E)** Corresponding TLC stained with orcinol to detect carbohydrate-containing species. A band at *Rf* ≈ 0.75 is enriched in the 100% MeOH fraction. **(F–H)** Representative flow cytometry histograms of lectin binding to Mm WT, Δ*pks15/1*, Δ*tesA* and capsule-stripped WT bacteria. Histograms show fluorescence intensity distributions for unlabeled control bacteria and bacteria incubated with ConA **(F)**, CF640R-labeled DscA **(G)** or CF640R-labeled DscE **(H)**. Dashed lines indicate the fluorescence peaks of unlabeled WT bacteria and lectin-labeled WT bacteria. Mm, *M. marinum*; ConA, concanavalin A; MPA, molybdophosphoric acid.

We therefore examined the polar-lipid fraction. Capsular and polar extracts shared an orcinol-positive band at *R*f ≈ 0.75 (Figure 6B), consistent with a glycolipid carrying the binding epitope. Its polarity and resistance to enzymatic and proteolytic treatments are compatible with a LOS-like glycolipid (Figure 5C).

To resolve the binding species further, Mmar polar lipids were subjected to two-step solid-phase extraction. The extract was first applied to a silica cartridge under highly aqueous conditions and eluted with increasing methanol concentrations. Binding activity was recovered mainly in the flow-through and initial water wash, whereas additional polar components remained on the matrix (Suppl. Figure S5B). These active fractions were pooled and separated on a C18 reversed-phase cartridge by stepwise methanol elution. Activity was largely absent from intermediate methanol fractions and from the final acetonitrile eluate but was recovered in the flow-through and wash and, more weakly, in the 100% methanol fraction (Figure 6C). Recovery under strong organic elution is consistent with an amphipathic glycoconjugate. TLC showed that the complex total extract was simplified in the 100% methanol fraction, in which a species at *Rf* ≈ 0.75 predominated; a band at this position was also present in the starting extract and in the other fractions. Positive staining with molybdophosphoric acid and orcinol showed a carbohydrate-containing lipid (Figure 6D,E). A weaker, lower-*R*f orcinol-positive species present across several fractions probably represented a minor co-purifying component. A species with closely matching migration was independently recovered by low-temperature acetone dissociation of DscA complexes, which precipitated the protein while preserving soluble lipid-associated material. The recovered fraction retained DscA-binding activity and matched the C18-enriched TLC profile, further linking binding to the glycolipid at *Rf* ≈ 0.75 (Suppl. Figure S5C).

These findings identify a dominant Proteinase K-resistant polar glycolipid as the main Mmar-binding species. Its identity remains unknown, but its biochemical behavior is consistent with a LOS-like molecule. Discoidins therefore mainly recognize a carbohydrate epitope on a non-proteinaceous polar surface glycolipid in Mmar, whereas binding to *M. bovis* BCG appears to involve a glycosylated proteinaceous capsular component.

### Discoidin binding is dependent on mycobacterial cell surface organization

Because the array identified PGL-associated ligands, we next tested whether disrupting PGL biosynthesis alters discoidin binding to the bacterial surface.

Deletion of *pks15/1*, which blocks PGL synthesis, mildly reduced DscA and DscE binding (approximately 20% and 10%, respectively). Deletion of *tesA*, which blocks PDIM and PGL synthesis, reduced binding much more strongly (approximately 85% for DscA and 45% for DscE) (Figure 6G,H), although PDIM is not a glycolipid. The Mmar Δ*tesA* mutant also showed increased ConA binding by approximately 20%, suggesting that loss of PDIM/PGL unmasks ConA ligands while reducing discoidin binding (Figure 6F). To distinguish ligand loss from altered surface availability, capsule and apolar-lipid fractions from WT, Δ*pks15/1* and Δ*tesA* Mmar were analyzed by dot blot. Binding remained comparable across strains, with no appreciable reduction in mutant extracts (Suppl. Figure S5D).

These observations support a direct contribution of PGL-associated glycans but also indicate additional ligands. PDIM/PGL loss may increase the accessibility or extraction of other ligands, including LOS-associated glycoconjugates, so binding is maintained in isolated fractions despite its reduction on intact bacteria.

Mechanical removal of the outer capsule likewise altered lectin binding (Figure 6F-H). ConA binding to stripped WT bacteria increased, indicating greater exposure of Man-LAM (Figure 6F). DscA binding decreased by approximately 50% (Figure 6G), whereas DscE binding increased slightly, by approximately 20% (Figure 6H). Thus, discoidin recognition depends on both glycolipid ligands, including PGL-associated structures, and their accessibility and spatial organization in the outer Mmar envelope.

### Various truncations or absence of LOS alter discoidin binding

Because LOS-associated saccharides were among the strongest array ligands, the transposon screen recovered several envelope-related hits, and biochemical fractionation identified a glycolipid with LOS-like properties, we tested whether LOS contribute to discoidin binding to the Mmar surface. A panel of mutants targeting different biosynthetic steps was compared with the corresponding complemented strains (Figure 7A). Disruption of MMAR_2349 (*wbbL2*) blocks early LOS glycosylation, producing highly truncated LOS-0/LOS-I intermediates. Mutations in MMAR_2331 and MMAR_2321 impair later maturation steps, allowing intermediate LOS-II/LOS-III forms but preventing fully elaborated LOS-IV. Insertional inactivation of *papA4* (MMAR_2343::tn) abolishes LOS biosynthesis and all detectable LOS species (Figure 7A).

**Figure 7.**
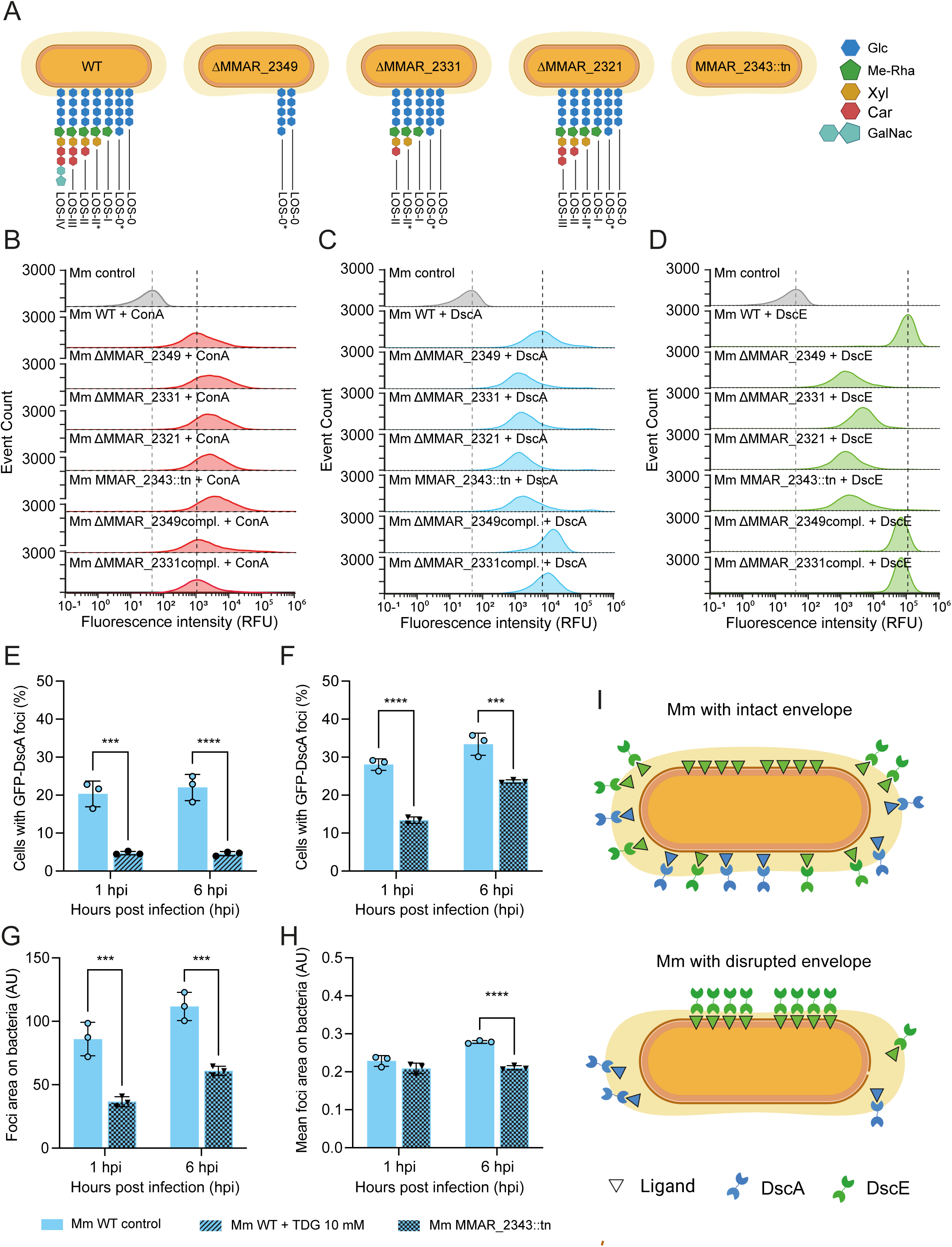
LOS-dependent glycan presentation controls discoidin binding and recruitment during infection. **(A)** Schematic representation of the LOS species produced by wild-type Mm and the LOS-biosynthesis mutants Δ*MMAR_2349*, Δ*MMAR_2331*, Δ*MMAR_2321* and *MMAR_2343*::tn. Monosaccharides are indicated as glucose (Glc), methylated rhamnose (Me-Rha), xylose (Xyl), caryophyllose (Car) and *N*-acetylgalactosamine (GalNAc). Asterisks denote aberrant LOS intermediates: LOS-II* lacks the terminal caryophyllose residue, whereas LOS-0* contains an additional glucose residue compared with LOS-0. **(B–D)** Representative flow cytometry histograms showing the binding of fluorescently labeled ConA **(B)**, DscA **(C)** and DscE **(D)** to wild-type Mm, the indicated LOS mutants and the corresponding complemented strains. Unlabeled bacteria are shown as controls. Vertical dashed lines indicate the reference fluorescence peaks of unlabeled bacteria and wild-type bacteria labeled with the corresponding lectin. **(E)** Quantification of GFP-DscA foci formation in *atg1*-knockout *D. discoideum* cells infected with wild-type Mm in the presence or absence of 10 mM TDG. The percentage of infected cells containing GFP-DscA foci was measured at 1 and 6 hpi. **(F–H)** Quantification of GFP-DscA recruitment in *atg1*-knockout *D. discoideum* cells infected with HADA-labeled wild-type Mm or the LOS-deficient *MMAR_2343*::tn mutant. The percentage of infected cells containing GFP-DscA foci **(F)**, the total area occupied by bacteria-associated GFP-DscA foci **(G)** and the mean area of individual bacteria-associated foci **(H)** were quantified at 1 and 6 hpi. **(I)** Model illustrating how LOS-dependent changes in envelope organization alter the accessibility and spatial distribution of surface glycan ligands, thereby modulating DscA and DscE recruitment. Data in **(E–H)** represent the mean ± SD of three independent experiments; individual experiments are shown. Statistical significance was assessed by two-way ANOVA followed by Šídák’s multiple-comparisons test. \*\*\**P* < 0.001; \*\*\*\**P* < 0.0001. Mm, *M. marinum*; LOS, lipooligosaccharide; TDG, thiodigalactoside; hpi, hours post-infection; AU, arbitrary units.

Flow cytometry showed a moderate increase in ConA binding across LOS-deficient strains, most prominently in the *papA4* transposon mutant; complementation reversed this phenotype (Figure 7B). In contrast, binding of both DscA and DscE to the bacterial surface was strongly reduced in mutants lacking the entire LOS repertoire or producing truncated LOS glycoforms (Figure 7C,D). DscA binding decreased by approximately 70% across the mutants, whereas DscE binding decreased by nearly 97% relative to WT bacteria. Complementation restored both signals to wild-type levels. Together, the glycan-array and genetic data show that LOS integrity is a major determinant of discoidin binding to Mmar, while the residual binding indicates that other envelope components also contribute.

### Absence of LOS reduces discoidin foci during infection of *D. discoideum*

Having established that discoidin binding to intact Mmar requires an intact H-type lectin pocket and is strongly influenced by LOS integrity, we next asked whether glycan-dependent binding also governs discoidin recruitment during intracellular infection. GFP-DscA was selected for this analysis because its robust foci signal during infection enabled reliable quantification of recruitment. Experiments were performed in *atg1*-knockout *D. discoideum*, in which loss of autophagy-related membrane repair accelerates Mmar escape from the MCV and thereby increases exposure of the bacterial surface to cytosolic lectins (17). The membrane-impermeant disaccharide TDG reduced infected cells harboring GFP-DscA foci by approximately 77% at 1 hpi and 79% at 6 hpi (Figure 7E), indicating that TDG reaches cytosolic discoidins after MCV damage. Infection with the LOS-deficient MMAR_2343::tn mutant produced a convergent phenotype: foci-positive cells decreased by approximately 52% at 1 hpi and 29% at 6 hpi (Figure 7F). Total bacteria-associated focus area decreased by approximately 57% and 45%, respectively (Figure 7G), whereas mean individual focus area was less affected, decreasing by approximately 9% and 24% (Figure 7H). Thus, blocking the host lectin pocket or removing a major bacterial ligand impaired intracellular recruitment. Residual recruitment to the LOS-deficient mutant indicates that other surface glycans also contribute. Discoidins therefore recognize multiple glycan epitopes through the β-galactoside-binding pocket *in vivo*, with LOS representing an important but not exclusive ligand in the Mmar envelope (Figure 7I).

## Discussion

The mycobacterial cell envelope is a highly stratified and dynamic structure in which distinct layers cooperate to define both the physical resilience of the bacillus and its immunological visibility to the host. A covalent AG–PG core anchors mycolic acids in the inner leaflet of the mycomembrane, whereas the outer leaflet contains non-covalently associated lipids and glycolipids, including trehalose-based lipids, PDIM, PGL and, in some species, LOS or GPLs. An external capsule contains polysaccharides, lipids, glycoproteins, GPLs and other envelope-derived material that can be remodeled or shed during growth and host interaction. This organization controls permeability and resistance, but also the accessibility of surface PAMPs to host sensors.

Our findings identify discoidins as cytosolic lectins that directly engage the mycobacterial surface and recognize a restricted set of carbohydrate determinants on lipid-associated glycoconjugates. DscA and DscE showed largely overlapping specificities across imaging, flow cytometry, fractionation and glycan array experiments (Table 1). Both bound intact Mmar through an intact H-type lectin pocket and preferentially recognized outer-envelope glycolipids rather than the major structural polysaccharides of the wall or capsule. All four canonical discoidins were recruited to damaged Mmar-containing compartments, suggesting that this selectivity extends beyond the two isoforms characterized biochemically. In contrast, discoidins were not recruited to endolysosomal compartments following LLOMe-induced sterile damage, showing that membrane damage alone is insufficient for their recruitment. DscA and DscE also bound HeLa but not growing *D. discoideum* cells, although WGA and HPA confirmed the presence of accessible amoebal glycans. Discoidins therefore appear to sense non-self glycans rather than membrane damage alone. DscA and DscE were nevertheless not equivalent. During infection, DscA formed foci on bacteria and at distal sites, whereas DscE remained mainly at the exposed bacterial surface (Table 1, row 1), in line with recognition of both bacilli and shed glycosylated envelope material that traffics within infected cells and is disseminated to bystander cells (49). DscE bound broth-grown Mmar more strongly, whereas cross-species FACS showed strongest DscA binding to *M. avium* and strongest DscE binding to Mmar (Table 1, rows 2–3). Pocket mutations and TDG abolished DscA binding and markedly reduced, but did not abolish, DscE binding; the two synthetic methylated L-rhamnose-containing disaccharides inhibited their respective discoidin isoform (Table 1, rows 4–5). This isoform specificity indicates chemically distinguishable pockets despite recognition of related carbohydrate determinants. DscA also recognized a broader glycan repertoire, while DscE was more selective for PGL- and LOS/GPL-associated epitopes; mechanical stripping decreased DscA but increased DscE binding (Table 1, rows 8 and 10). These differences support distinct epitope preferences between their H-type lectin pockets despite shared Gal/GalNAc specificity (33, 34, 47). Because dominant mycobacterial envelope glycans differ from previously described discoidin ligands, including β-GalNAc-containing structures (34), additional bacterial glycoconjugates likely contribute during infection.

**Table 1.**
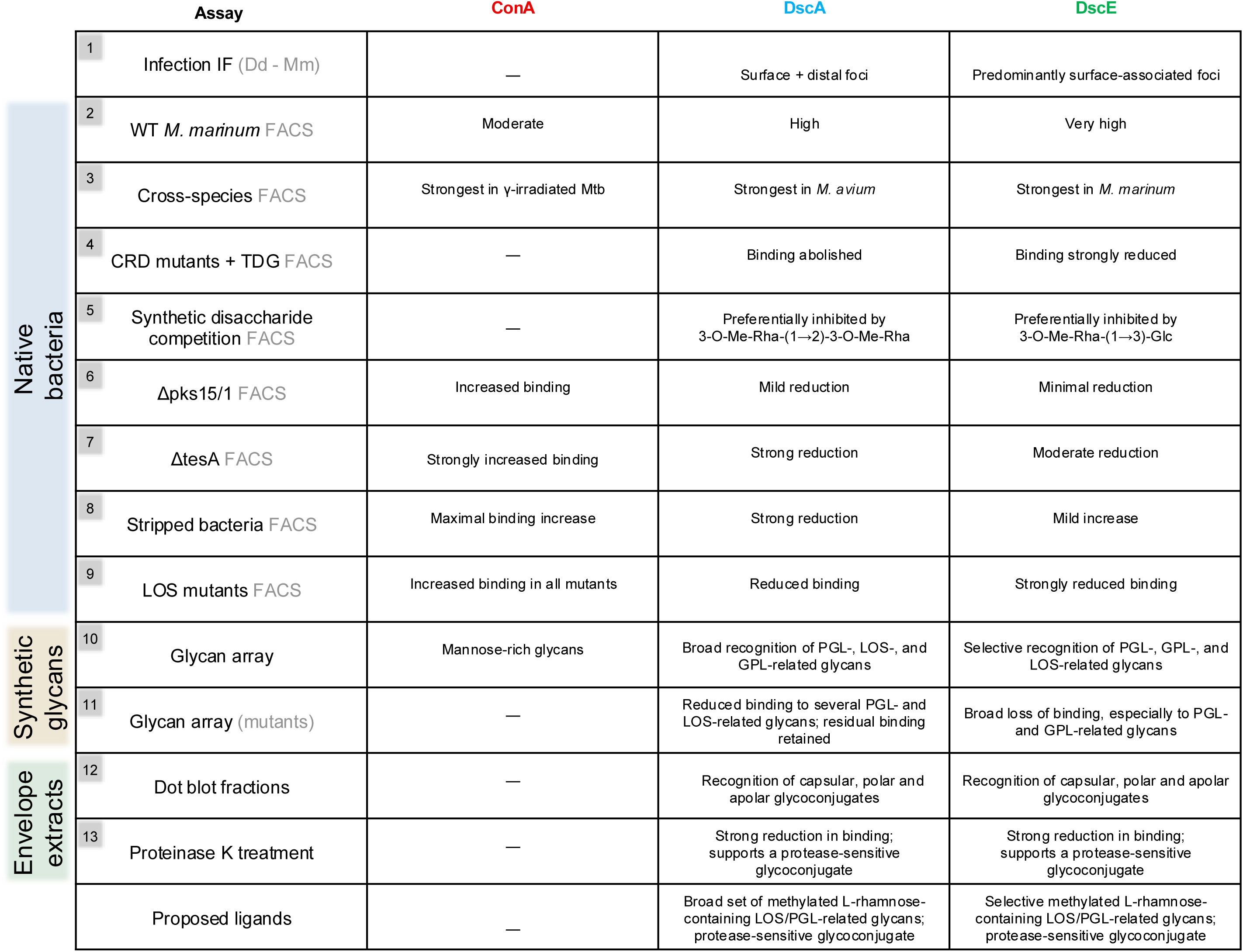
Integrated summary of discoidin recognition across experimental assays. Comparison of ConA, DscA and DscE bindings monitored by a variety of assays performed with either native bacteria (FACS), or biochemical fractions of the envelope (dot blots) or a synthetic glycan array, using Mmar wt and cell-wall mutants. Together, the data support methylated L-rhamnose-containing glycan epitopes harbored by a variety of lipid-associated glycoconjugates, including LOSs, PGLs and GPLs, as major discoidin ligands in Mmar. The table further illustrates how envelope organization and ligand accessibility shape discoidin recognition at the bacterial surface.

The best-supported ligands are a restricted set of outer-envelope carbohydrate epitopes, including methylated L-rhamnose motifs. Dot blots detected binding in capsular, polar and apolar fractions, with the strongest signals in capsular and polar extracts (Table 1, row 12), but excluded AG– PG, AM/LAM, α-glucan and purified TDM as major ligands. The synthetic array likewise identified a narrow set of PGL-, LOS- and GPL-associated glycans, while mannose-rich LAM structures and capsular α-glucans were not significantly recognized (Table 1, row 10). Because the array reports carbohydrate specificity without identifying the native carrier, binding to a GPL-derived glycan indicates a shared epitope but does not establish GPLs as ligands in Mmar or *M. bovis* BCG, neither of which produces canonical GPLs (50–52). Fractionation of Mmar also enriched a carbohydrate-containing amphipathic lipid that co-fractionated with binding activity (Table 1, row 12). Its identity remains unresolved, but its biochemical properties, considered together with the array and mutant data, are consistent with LOS-containing glycolipids. Proteinase K sensitivity further indicated species-specific carrier classes. Because canonical GPL peptide cores are expected to resist standard proteases, complete loss of binding to *M. bovis* BCG extracts instead suggests that most epitopes occur on proteins or glycoconjugates with protease-sensitive peptide bonds, whereas Mmar retained a protease-resistant fraction consistent with LOS or other glycolipids (Table 1, row 13). Thus, discoidin-binding epitopes can be presented by distinct carrier classes in different mycobacterial species. Ligand presence and accessibility were not equivalent. Methylated L-rhamnose-containing glycolipids remained detectable in extracts from several Mmar mutants, although binding to intact bacilli was reduced. This was clearest for PDIM/PGL mutants (Table 1, rows 6–7): deletion of *pks15/1*, which affects PGL maturation, caused a modest reduction, whereas deletion of *tesA*, which abolishes PDIM and PGL synthesis, caused a much greater loss of surface binding despite preserved binding to isolated fractions. This uncoupling separates biochemical ligand abundance from native surface display. This residual activity is consistent with other methylated L-rhamnose-containing glycoconjugates retaining lectin binding when mature PGL is absent. Discoidin recognition therefore depends on the native presentation of several related glycoconjugates, not simply their presence in the envelope.

The transposon-screen hits also implicated outer-envelope organization more than individual glycan-biosynthetic enzymes. They included PDIM/PGL biosynthetic genes (*fadD2*, *fadD28* and *mas*), Type VII/ESX substrates that shape the capsule (including PE, PPE and PE-PGRS proteins), and other, largely uncharacterized membrane-associated proteins (45). ESX-5-dependent PPE secretion controls capsular integrity and surface hydrophobicity (45, 53); disrupting these genes may therefore alter glycolipid display without eliminating a specific ligand. Comparison with their H37Rv counterparts revealed no convergence on a known regulon (e.g., DosR, KstR/KstR2, *phoP*, IdeR or Zur), suggesting that the mutations affect multiple envelope-organization pathways and do not represent a shared regulatory process. LOS mutants provided further support for this model. Mutations affecting different maturation steps reduced DscA and DscE binding, and complementation restored near-wild-type levels. The shared phenotype argues against exclusive recognition of a single terminal epitope: discoidins may recognize multiple LOS glycoforms, while LOS-dependent envelope organization may also regulate the exposure of other glycan determinants. This dual role is consistent with evidence that LOS perturbation affects colony morphology, membrane properties and export of surface material (54, 55). LOS may therefore contribute directly as a ligand and indirectly by organizing the outer envelope.

Complementary perturbations of the host lectin and bacterial ligand established the relevance of this interaction during infection. Both TDG-mediated inhibition of the DscA lectin pocket and bacterial LOS deficiency reduced the proportion of infected cells containing GFP-DscA foci. These convergent effects link glycan recognition to intracellular recruitment and identify LOS-dependent glycan presentation as an important, but non-exclusive determinant. Residual recruitment to the LOS-deficient mutant supports the contribution of additional surface glycans *in vivo*. Mechanical stripping further showed that surface organization controls lectin access: ConA binding increased, consistent with unmasking of mannose-containing glycoconjugates, while DscA binding decreased and DscE binding increased modestly. Similar context dependence governs DC-SIGN recognition of surface-exposed Man-LAM (20, 22, 23). Discoidins also parallel metazoan galectins as soluble cytosolic lectins that undergo noncanonical secretion and accumulate at damaged pathogen-containing compartments. However, intracellular galectins primarily detect host self glycans exposed by membrane damage (24, 39, 56), whereas our data indicate that discoidins directly recognize exposed microbial glycoconjugates. This contrast raises the possibility that discoidins represent an ancestral form of cytosolic lectin surveillance based on direct recognition of specific PAMP, whereas galectins provide a more general system that detects self-glycan DAMPs exposed by infection-induced or sterile membrane damage.

Overall, discoidins act as cytosolic probes of glycoconjugate exposure at the mycobacterial surface. They recognize overlapping but non-equivalent sets of lipid-associated glycans enriched in methylated L-rhamnose, including LOS-, PGL- and GPL-related epitopes, whose accessibility is controlled by outer-envelope organization.

## Materials and Methods

### *Dictyostelium discoideum* strains and culture conditions

*D. discoideum* was cultured axenically at 22 °C in HL5c (Formedium) with penicillin/streptomycin (100 U/mL/100 μg/mL). GFP-discoidin transformants were obtained after electroporation and selection with hygromycin (15 μg/mL, knockouts; 50 μg/mL, act5 integrants) or blasticidin (5 μg/mL, KI/KO lines).

### Mycobacterial strains and culture conditions

Mmar/*M. smegmatis* grew at 32 °C; *M. brumae*, *M. abscessus*, *M. avium*, *M. bovis* BCG, and *M. kansasii* at 37 °C, in Middlebrook 7H9 (Difco)/10% OADC/0.2% glycerol/0.05% Tween-80 or tyloxapol, at 150 rpm with 5-mm glass beads. Selection used hygromycin (100 μg/mL) or kanamycin (25–50 μg/mL); Δpks15/1, ΔtesA, LOS, and complemented strains used identical conditions. Gamma-irradiated *M. tuberculosis* H37Rv cells (NR-49098) came from BEI Resources.

### Infection assays and live-cell imaging

*D. discoideum* was infected with Mmar (MOI 10); uptake was synchronized by two centrifugations (1,000 × g, 10 min) and 10–20 min at 25 °C before washing. Cells expressing GFP-discoidins were imaged live on a Leica SP8/Stellaris 8 (63×/1.4 NA). Cells were also fixed using ultra-cold methanol or 4% PFA and processed for immunofluorescence. Discoidins were detected with a rabbit polyclonal anti-discoidin antibody (7-78-3; 1:500; gift of W. Nellen), followed by a goat anti-rabbit AlexaFluor647-conjugated antibody (1:1,000; Molecular Probes).

For LOS experiments, wild-type and LOS-deficient Mmar were pre-labeled with HADA (1 h) and infected cells were fixed with 4% PFA at 1 and 6 hpi. For TDG experiments, cells were pre-treated overnight with 10 mM TDG, maintained during infection, and imaged live at 1 and 6 hpi versus untreated infected controls. Imaging was performed by automated high-content (HC) microscopy.

### LLOMe-induced endolysosomal damage

GFP-Vps32/DscA/DscE expressing cells were treated with 5 mM LLOMe and imaged by time-lapse microscopy for 15 min on a Nikon Ti/CSU-W1 spinning disk (100×; identical settings); images at 0 and 10 min are shown in Figure 1B. Untreated cells served as negative control and GFP-Vps32 recruitment as a positive control for damage.

### Labeling of bacterial surface glycoconjugates and peptidoglycan

Surface glycoconjugates were oxidized in 1 mM periodate/0.1 M acetate (pH 5.5; 20 min, 4 °C), coupled to 1 mM AlexaFluor488 (AF488) hydrazide (1 h, room temperature), glycerol-quenched, and washed. Mmar peptidoglycan was labeled for 1 h at 32 °C with 500 μM HADA [3-[[(7-hydroxy-2-oxo-2H-1-benzopyran-3-yl)carbonyl]amino]-D-alanine hydrochloride] followed by PBS washes.

### Expression, purification, and labeling of recombinant discoidins

Codon-optimized DscA/DscE and lectin-pocket mutants were cloned in pProExHTb as N-terminal His₆ fusions, expressed in *E. coli* Rosetta2(DE3) pLysS, and induced overnight (0.2 mM IPTG, 16 °C). Ni²⁺ HisTrap HP-purified proteins were buffer-exchanged, 10-kDa-MWCO-concentrated, NHS-labeled with CF640R or AF532, and stored in PBS/0.5 mM CaCl₂/1% BSA.

### In vitro lectin binding assays

Bacteria (1 × 10⁷) were incubated with labeled discoidin (100–600 μg/mL; 1 h, room temperature) in PBS/0.5 mM CaCl₂/1% BSA, then washed and imaged or analyzed by Sony SH800 flow cytometry; CF640R-concanavalin A (Biotium) was used as a control lectin. For competition, 10 mM TDG (Cayman) in PBS or 10 mM of each synthetic disaccharide (dissolved in methanol; 2.5% v/v final) was added. HeLa and *D. discoideum* cells were fixed with 4% PFA and incubated with CF640R-DscA/DscE, AF488-WGA or AF488-*Helix pomatia* agglutinin (HPA) (10–50 μg/mL), then washed and imaged.

### FACS-based enrichment of DscE low-binding mutants

A published Mmar transposon library (57) was incubated with fluorescent DscE; the lowest-5% fluorescent Mmar were FACS-isolated, expanded in 7H9, and treatment repeated twice before genomic-DNA extraction and insertion sequencing.

### Data analysis of transposon insertion sequences

Cutadapt (-e 0.05 --front CGGGGACTTATCAGCCAACCTGT --minimum-length 25 --maximum-length 85) was used to retain reads bearing the transposon sequence before TA sites. Trimmed reads were mapped on Mmar M RefSeq assembly GCF_000018345.1 with HiSAT2 (--no-spliced-alignment) and the number of insertions was quantified at every TA site.

### Bioinformatic analysis of genes with Tn insertions

Mmar M ORFs (UniProtKB UP000001190) (60) used InterProScan 5 (61)/Pfam (62), SignalP (63), TMHMM (64), Phobius (65), and tRNAscan-SE (66). *M. tuberculosis* H37Rv orthologues used reciprocal Smith–Waterman matches (BLOSUM62; gap-open/extension 11/1) via parasail (67)/Biopython (68) and independent progressiveMauve synteny (69); reciprocal/one-way matches were high-confidence/tentative. High-confidence hits were compared with DosR, KstR/KstR2, PhoP, IdeR, and Zur regulons and type-VII toxin loci (70). Ambiguous hits used Foldseek (74) to compare AlphaFold models (71)(72) with PDB (73), plus PaperBLAST (75).

### Structural modeling and visualization

DscA/DscE crystals (PDB 2W95/2VMC) and AlphaFold 3 DscC/DscD predictions were retrieved. DscC/DscD homotrimers used DscA-trimer geometry; assembly, visualization, domain coloring, and figures used UCSF ChimeraX 1.11.1.

### Cell lysis and recovery of the mAGP complex

Cell pellets washed in 200 mL PBS were treated with DNase I (10 mg/mL; 30 min, ice), probe-sonicated, heated in 1% (w/v) SDS (115 °C, 15 min), and centrifuged (12,000 × g, 1 h). Pellets were washed repeatedly in 150 mL Milli-Q water at 25 °C until SDS-free and lyophilized.

### Isolation of arabinogalactan–peptidoglycan

Lyophilized material was stirred in 60 mL 0.5% (w/v) KOH/methanol (37 °C, 4 days), cooled to 4 °C, and centrifuged (1,200 × g, 1 h). Pellets were washed twice in ice-cold methanol and four times in diethyl ether, then stirred overnight at 55 °C in 60 mL PBS/15 μg/mL proteinase K. AGP was recovered (27,000 × g, 30 min, 25 °C), washed twice, snap-frozen, and lyophilized.

### Purification of peptidoglycan

To separate soluble arabinogalactan from peptidoglycan, AGP was hydrolyzed in 0.2 M H₂SO₄ (2 h, 60 °C) and centrifuged (12,000 × g, 20 min, 20 °C). The pellet was neutralized in 0.5 M Na₂CO₃, washed thrice, and digested (8 h, 37 °C) with α-amylase/DNase I/RNase A (100/10/5 μg/mL), then overnight with proteinase K (100 μg/mL). It was refluxed 3 h in 1% SDS to remove leftover lipids and proteins, washed seven times (12,000 × g, 20 min, 20 °C), flash-frozen, and lyophilized.

### Capsule extraction

Capsules from Mmar M, *M. bovis* BCG Danish 1331, and its *lamH::Himar1* mutant (58) were extracted identically: confluent 7H10 agar cultures were scraped, water-suspended, and vortexed (1 min, 1,200 rpm). After pelleting (2,000 × g, 10 min), supernatants were 0.45-μm-filtered, frozen, lyophilized, and redissolved in ddH₂O (1 mg/mL).

### Bligh–Dyer lipid extraction

Bligh–Dyer extraction (59) used 1 mL samples + 2.0 mL methanol (vortex 30 s) + 1.0 mL chloroform (vortex 30 s; equilibrate 10 min; CHCl₃:MeOH:H₂O 1:2:1). After adding 1.0 mL chloroform/0.8 mL water, samples were shaken 15 s and centrifuged (1,500 × g, 5 min). The lower phase and one 1.0-mL-chloroform re-extract were pooled and dried.

### Purification of lipoarabinomannan and lipomannan

Cultures were pelleted (6,000 × g, 10 min), PBS-washed twice, suspended in 10 mL PBS, and bead-beaten (3 × 45 s, 6.0 m/s). Lysates received two equal-volume phenol extractions (85 °C, 2 h; vortex every 30 min); the centrifuged phenol phase was re-extracted with equal-volume water. Pooled aqueous phases were dialyzed (3.5-kDa MWCO) against six 1-L water changes (3 h each, room temperature), lyophilized, and stored at −20 °C.

### Dot Blot Binding Assay

Aqueous/solvent extracts were spotted (2 μL; 1-cm intervals) on 2 × 7-cm nitrocellulose/PVDF strips, respectively, and dried 2 h. Strips were blocked in PBS/3% (w/v) BSA/0.5 mM CaCl₂ (1 h, 20 °C, gentle agitation), then incubated in 5 mL buffer with 1 μg/mL AF532- or CF640R-DscA/DscE (1 h, 20 °C, 300 rpm). Three 15-min washes and 2 h air-drying preceded Typhoon scanning at label-specific wavelengths.

### Enzymatic and proteolytic treatments

Capsules were treated with α-amylase/pullulanase [α-(1→4)/(1→6) glucosidic bonds] or proteinase K (1–5 mg/mL, 16 h), then heat inactivated, dialyzed, and used for dot blot.

### Thin-layer chromatography (TLC)

10 μL of polar/apolar lipids (1 mg/mL in methanol/chloroform) or silica/C18 fractions in eluent was spotted (in five 2-μL loads per lane dried between applications) on Merck Silica Gel 60 F₂₅₄ plates. Development used chloroform:methanol:water (60:30:6 or 30:8:1) or chloroform:methanol (95:5); air-dried plates were stained with 0.2% orcinol/concentrated H₂SO₄ or 10% molybdophosphoric acid/ethanol and heated at 110 °C.

### Silica and C18 fractionation of polar lipids

Polar lipids (1 mg/1 mL methanol) were N₂-dried, redissolved in 1 mL ddH₂O, and loaded on an Isolute Si cartridge (5-g bed/10-mL reservoir; Biotage) conditioned by gravity (∼1 mL/min) with 10 mL methanol then 50 mL ddH₂O. FT and 5-mL water wash W₀ were collected; 5-mL eluates used 5, 10, 15, 20, 30, 40, 60, 80, and 100% methanol, then 100% acetonitrile. Fractions were SpeedVac-dried (∼2 h, room temperature) and redissolved in 1 mL sterile ddH₂O.

FT + W₀ from the silica column were redissolved in 1 mL water and syringe-loaded on a Strata C18-E cartridge equilibrated by gravity (∼1 mL/min) with 50 mL HPLC methanol then 50 mL sterile water. C18-FT and 5-mL wash C18-W₀ were collected; 5-mL eluates used 5, 10, 25, 50, and 100% methanol (C18-E₁–E₅), then 100% acetonitrile (C18-E₆). All fractions were SpeedVac-dried at room temperature and redissolved in 1 mL sterile water.

### Pull down assays

Sonicated insoluble cell wall substrates (10 mg/mL PBS) were mixed 1 h at room temperature with 0.25 μM WGA or 1.0 μM BSA/DscA/DscE in PBS/0.5 mM CaCl₂ and washed with the same buffer. Input and two post-centrifugation supernatants were read at fluorophore-specific wavelengths; bound (%) = [input – (unbound + washes)]/input × 100.

Purified protein (100 μL, 8 mg/mL) plus dry-mass-normalized *M. marinum* polar lipids (1 mg/mL) was incubated 1 h at 20 °C. Ni-NTA-captured protein was washed twice in 20 mM Tris-HCl (pH 7.5)/150 mM NaCl and eluted with 500 mM imidazole in that buffer; Bligh–Dyer-extracted lipids were dried and TLC-analyzed. AG–PG was incubated with labeled discoidin (1 μM) in PBS/0.5 mM CaCl₂ 1 h at room temperature and centrifuged (13,000 × g). Supernatant depletion after two pellet washes quantified binding; WGA/BSA were positive/negative controls.

### Glycan microarray

Labeled DscA and DscE were used to probe Semiotik glycan array slides. The protocols and analysis pipeline are described in detail in the Appendix.

### Mycobacteria Glycan microarray

Labeled DscA and DscE were also used to probe mycobacteria-specific glycan array slides (47). The protocols and analysis pipeline are described in detail in the Appendix.

### Synthesis of disaccharide ligands

Complete details on the synthesis of the mycobacteria-related disaccharides evaluated as DscA and DscE ligands are found in the Appendix.

### Image analysis and statistics

Microscopy images were processed using Fiji, and quantitative analysis was performed using MetaXpress. Flow cytometry data were analyzed using Floreada.io. Statistical analyses were conducted in GraphPad Prism. Data are presented as mean ± SD from ≥3 independent experiments. Statistical significance was determined using appropriate tests as indicated in figure legends.

## Data availability

All data underlying the findings described in this manuscript are provided within the article and its supporting Information files and Appendix.

## Authors’ contributions

Conceptualization: DD, TS, GG, PM, TLL, MF

Methodology: DD, AF, PM, GG, TLL, LSM, MF, RBZ, GHH, SvG, LSL, CKC

Investigation: DD, TS, MF, TLL, PM, JP

Formal Analysis: DD, TS, MF, TLL, PM, JP

Data Curation: DD, TS, MF, GG, PM, TLL, LSM, GHH, SvG, LSL, CKC

Visualization: DD, TS, GG, PM, TLL, MF

Resources: PM, TLL, TS

Funding Acquisition: TS, PM, TLL

Project Administration: TS, DD

Supervision: TS, DD, MF, PM, TLL

Writing – Original Draft: DD, TS

Writing – Review & Editing: DD, TS, MF, GG, TLL, PM

## Funding

This work was supported by the Swiss National Science Foundation (grants 310030_188813 and 310030_219364), by Academia Sinica Grant AS-IDR-110-06, and an EMBO post-doctoral fellowship earned by MF (ALTF 715-2021). SvG received grant support by the Swiss National Science Foundation (Grants #320030_219451 and #310030_219605 (S.v.G.), and Swiss Cancer Research (KFS-5936-08-2023-R).

## Acknowledgments

We thank Dr Ana Teresa López Jiménez for performing preliminary insightful experiments during her thesis work. We thank the ACCESS facility, especially Dimitri Moreau and Stefania Vossio, for technical support and assistance. We are grateful to Laurent Kremer for providing the Mmar LOS mutant strains, and to Graham Stewart, Tom Mendum, and Rachel Butler for their help in generating the Mmar transposon libraries. Dr. Ronelito J. Perez is thanked for assistance on the glycan array experiments. We also acknowledge the protein purification and centrifuge core in the Institute of Biological Chemistry at Academia Sinica, the IBC NMR facility and the Academia Sinica High-Field NMR Center, funded by Academia Sinica Core Facility and Innovative Instrument Project (AS-CFII-111-214). Schemes in figures were partially created in BioRender, under CC BY 4.0 licence.

## Supplementary Figure and table legends

**Supplementary Figure S1.**
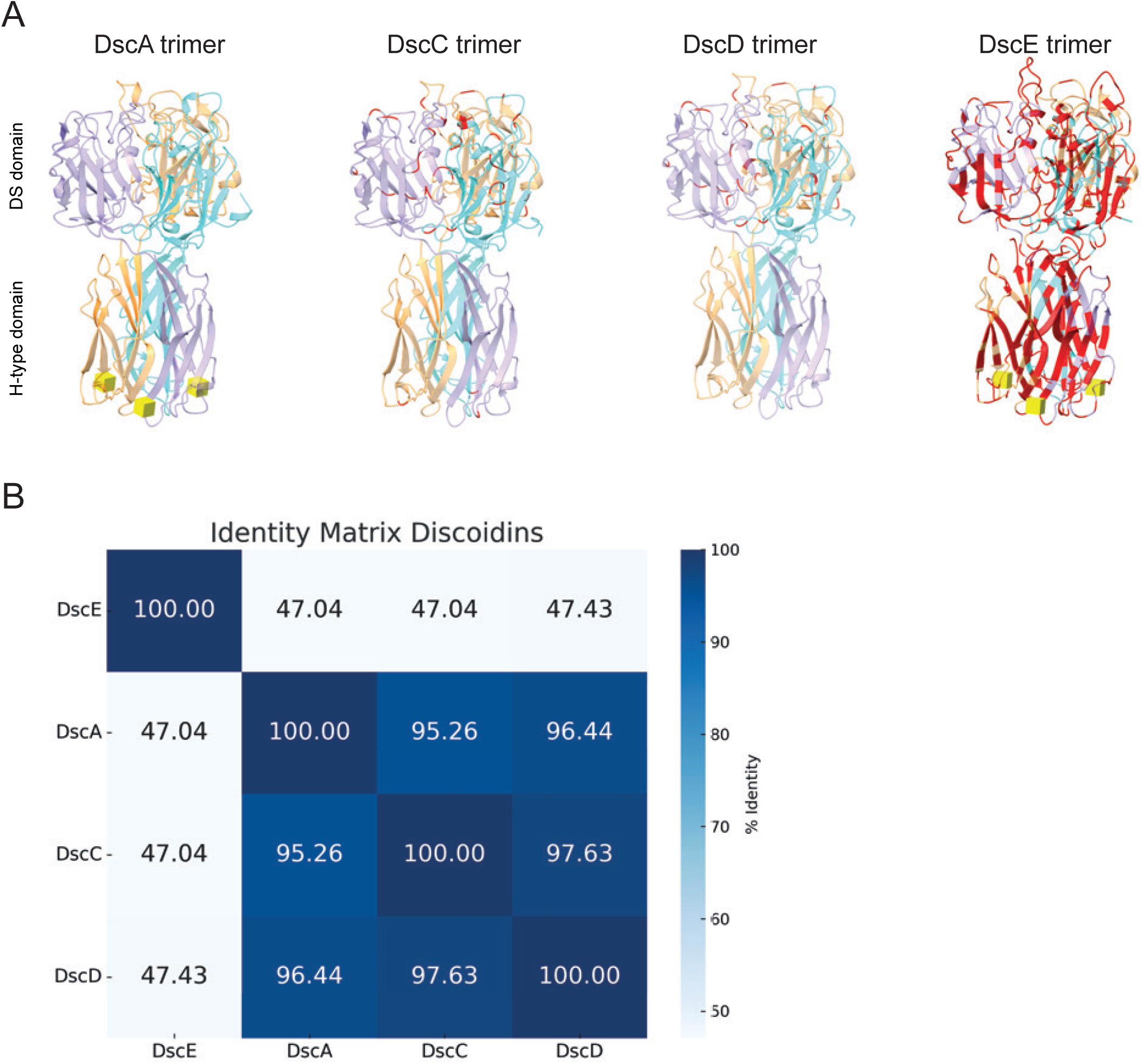
Structural comparison and sequence identity of canonical discoidins. **(A)** Structural representations of the DscA, DscC, DscD and DscE homotrimers. Experimentally determined crystal structures from the Protein Data Bank were used for DscA and DscE. Homotrimeric models of DscC and DscD were generated by assembling AlphaFold-predicted monomers according to the trimeric organization of DscA. Individual monomers are shown in cyan, orange and purple. Amino acid residues that differ from the corresponding positions in DscA are highlighted in red. The N-terminal discoidin (DS) domain and C-terminal H-type lectin domain are indicated. β-*N*-acetylgalactosamine (β-GalNAc) molecules bound within the H-type lectin pockets of DscA and DscE are shown in yellow. **(B)** Pairwise amino acid sequence identities among DscA, DscC, DscD and DscE were calculated with EMBOSS Needle (global alignment). Percentage sequence identities are indicated within the matrix and represented by the color scale.

**Supplementary Figure S2.**
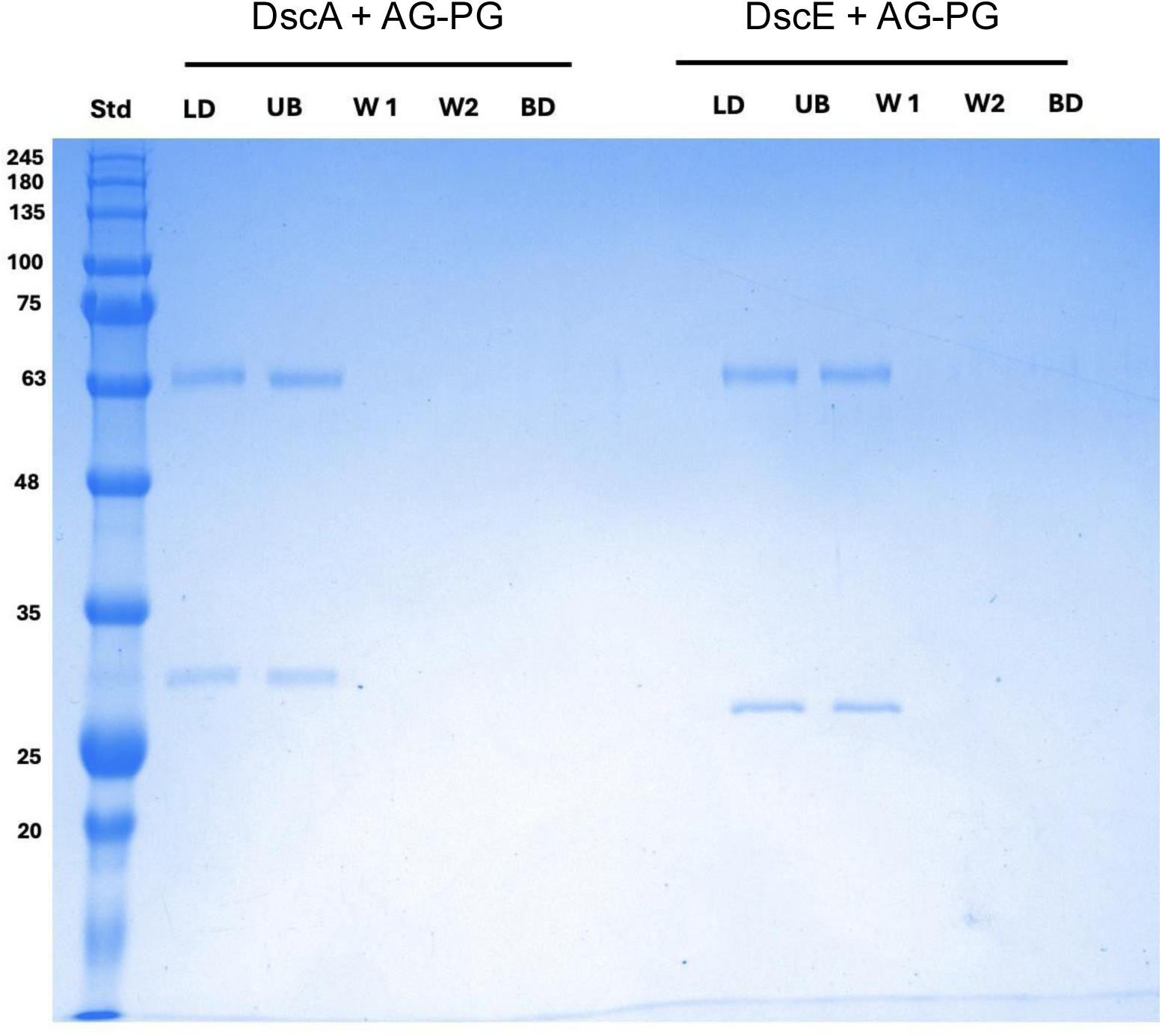
Discoidins do not bind purified AG–PG. Pull-down assays were performed by incubating recombinant DscA or DscE with purified arabinogalactan–peptidoglycan (AG–PG) from *M. marinum*. Following incubation, samples were separated into load (LD), unbound (UB), two wash fractions (W1 and W2) and bound fraction (BD). Proteins were analyzed by SDS–PAGE followed by Coomassie staining. DscA and DscE were detected in the load and unbound fractions but were not detectable in the wash or bound fractions, indicating no stable interaction with purified AG–PG under the conditions tested. Molecular mass markers, expressed in (kDa), are indicated on the left.

**Supplementary Figure S3.**
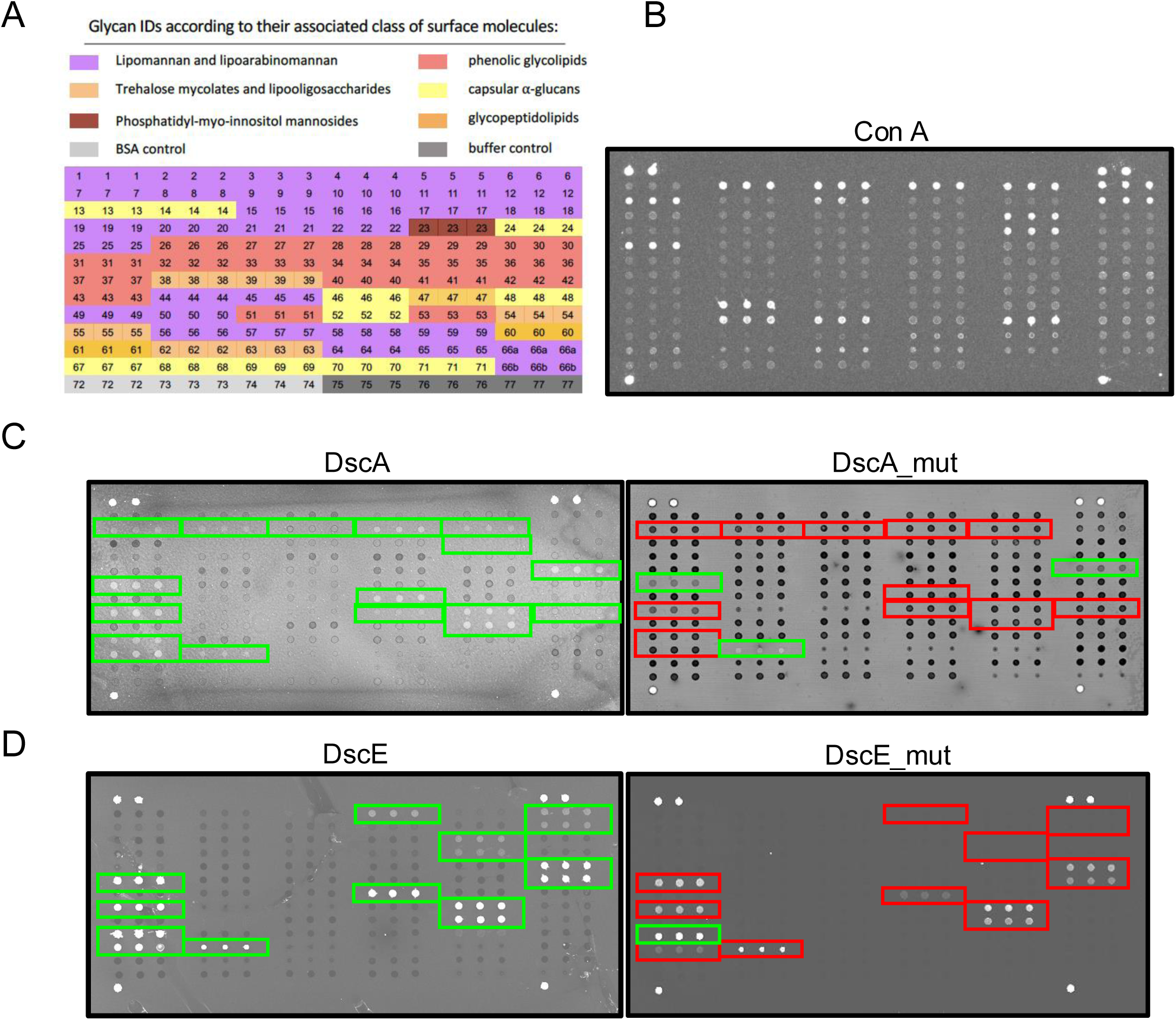
Glycan array profiling of wild-type and lectin-pocket mutant discoidins. **(A)** Layout of the synthetic mycobacterial glycan array, showing glycan IDs grouped according to their associated surface-molecule classes: lipomannan and lipoarabinomannan, trehalose mycolates and lipooligosaccharides, phosphatidyl-*myo*-inositol mannosides, capsular α-glucans, phenolic glycolipids and glycopeptidolipids. BSA and buffer control spots are also indicated. Each glycan was printed in triplicate. **(B)** Representative binding profile obtained with ConA, used as a positive-control lectin. **(C, D)** Representative array images showing the binding profiles of wild-type DscA and its lectin-pocket mutant DscA_mut (W235F/Y241S) **(C)**, and wild-type DscE and its lectin-pocket mutant DscE_mut (W238F/Y244S) **(D)**. Green boxes in the wild-type panels indicate recognized glycans. In the mutant panels, red boxes identify corresponding regions in which binding was reduced or lost, whereas green boxes indicate signals retained by the mutant proteins. Arrays were probed with fluorescently labeled recombinant proteins. ConA, concanavalin A; BSA, bovine serum albumin.

**Supplementary Figure S4.**
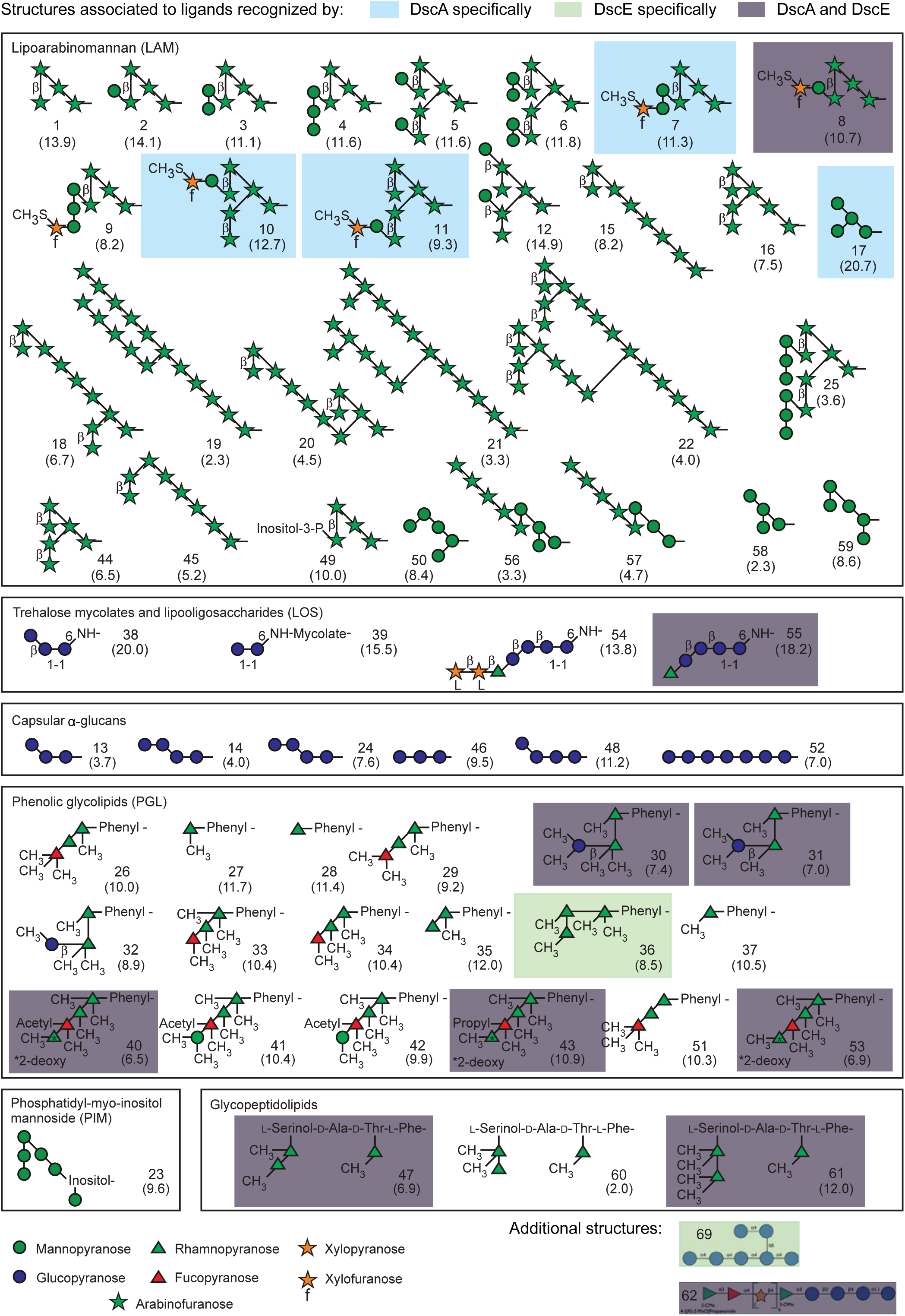
Mapping of glycan structures recognized by DscA and DscE on the synthetic mycobacterial glycan array. Annotated overview of glycan structures presented on the synthetic mycobacterial glycan array, organized according to their associated envelope molecule classes, including AM/LAM, trehalose mycolates and LOSs, capsular α-glucans, PGLs, PIMs, and GPLs. Colored overlays indicate glycans preferentially recognized by fluorescently labeled DscA (blue), DscE (green), or shared between both discoidins (gray), based on array binding analyses. Glycan structures were adapted from (47).

**Supplementary Figure S5.**
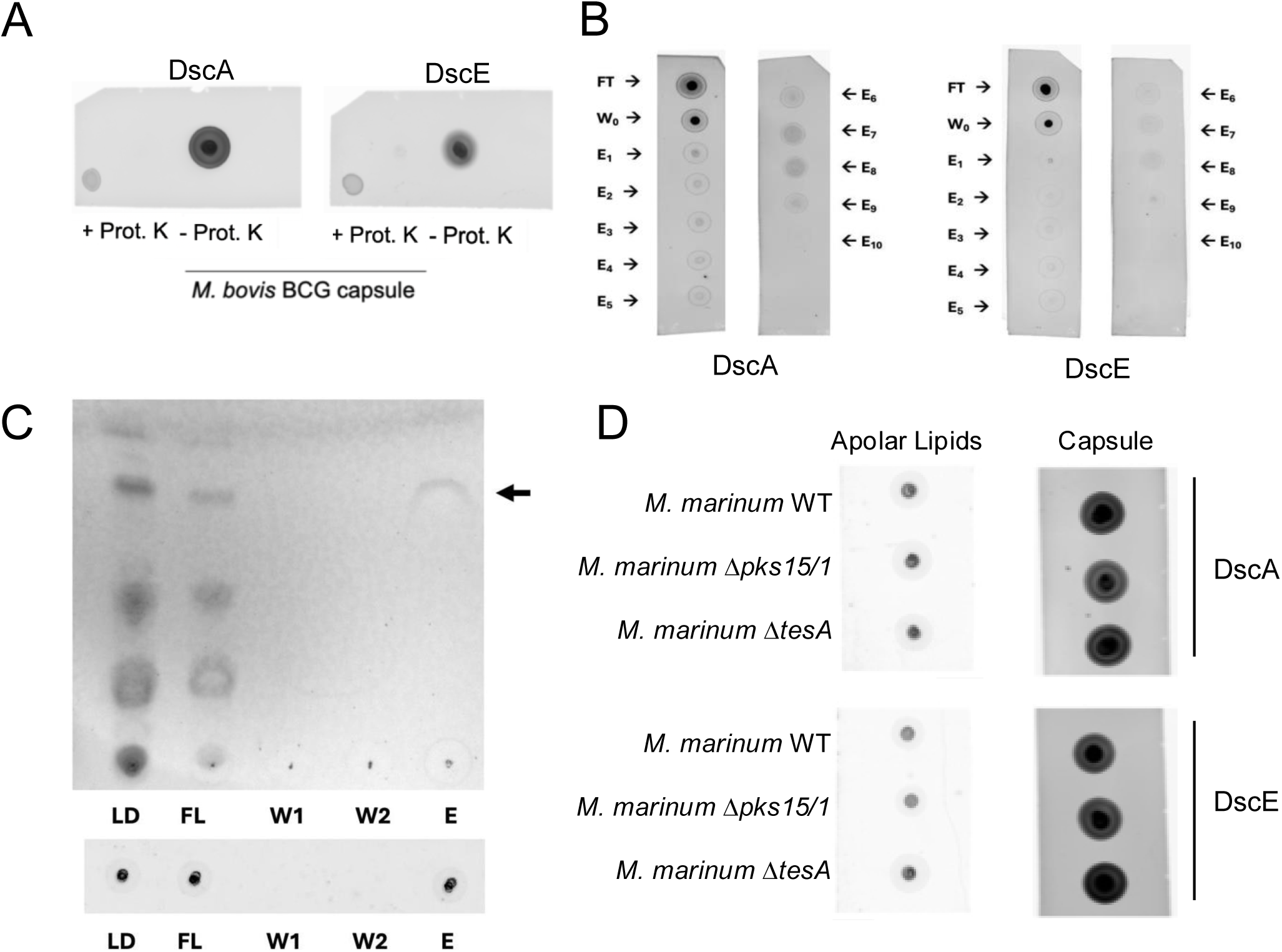
Biochemical characterization of mycobacterial envelope glycoconjugates carrying glycan epitopes recognized by discoidins. **(A)** Dot blot analysis of *M. bovis* BCG capsular extracts treated or not with proteinase K and subsequently probed with fluorescently labeled recombinant DscA or DscE. **(B)** Dot blot analysis of fractions obtained by silica solid-phase extraction of Mmar polar lipid extracts. The flow-through (FT), initial wash (W0) and sequential elution fractions (E1–E10) were probed with fluorescently labeled recombinant DscA or DscE. Material recognized by both discoidins was detected predominantly in the FT and W0 fractions. **(C)** Thin-layer chromatography analysis of fractions obtained from DscA pull-down and acetone-mediated ligand-release experiments. An orcinol-positive band recovered in the elution fraction (black arrow) migrated at an R*f* similar to that of the putative ligand previously identified in Mmar polar lipid fractions. The lower panel shows the corresponding dot blot, confirming retention of DscA-binding activity in the released fraction. LD, load; FL, flow-through; W1 and W2, wash fractions; E, elution fraction. **(D)** Dot blot analysis of apolar lipid and capsular fractions isolated from wild-type, Δ*pks15/1* and Δ*tesA* Mmar and probed with fluorescently labeled recombinant DscA or DscE. Mmar, *M. marinum*.

**Supplementary Table S1. List and classification of genes enriched in the DscE low-binding population. (A)** Table of genes enriched across the three rounds of selection in the transposon screen. Gene IDs, corresponding locus tags, gene names (when annotated), and number of transposon insertion sites are indicated. Genes are grouped by functional categories, including cell wall synthesis, cell wall-associated processes, central metabolism, DNA/RNA-related functions, respiratory chain, and two-component regulatory systems.

## Notes

### Competing Interest Statement

The authors have declared no competing interest.

### Summary of Updates

The manuscript was revised by incorporating important new data, including the use of synthetic discoidin ligands as binding competitors, use of additional eukaryotic and bacterial glycan arrays, as well as in vivo formation of discoidin foci in cells infected with M. marinum LOS mutants.

